# New Insights into the Genetic Diversity of the Bacterial Plant Pathogen ‘*Candidatus* Liberibacter solanacearum’ as Revealed by a New Multilocus Sequence Analysis Scheme

**DOI:** 10.1101/623405

**Authors:** Ahmed Hajri, Pascaline Cousseau-Suhard, Pascal Gentit, Marianne Loiseau

## Abstract

‘*Candidatus* Liberibacter solanacearum’ (Lso) has emerged as a serious threat on solanaceous and apiaceous crops worldwide. Five Lso haplotypes (LsoA, LsoB, LsoC, LsoD and LsoE) have been identified so far. To decipher genetic relationships between Lso strains, a MLSA study of seven housekeeping genes (*acnA, atpD, ftsZ, glnA, glyA, gnd* and *groEL*) was performed on a representative bacterial collection of 49 Lso strains. In all, 5415 bp spanning the seven loci were obtained from each of the 49 strains of our bacterial collection. Analysis of sequence data was consistent with a clonal population structure with no evidence of recombination. Phylogenies reconstructed from individual genes, and with concatenated data, were globally congruent with each other. In addition to the five highly supported and distinct genetic clusters, which correspond to the five established haplotypes, our phylogenetic data revealed the presence of a sixth haplotype, designated ‘LsoG’. This new haplotype is currently represented by two strains from France which had distinct sequences in four out of the seven tested housekeeping genes. Altogether, the data presented here provide new information regarding the genetic structure of Lso and the evolutionary history of the haplotypes defined within this bacterial species.

## Introduction

‘*Candidatus* Liberibacter solanacearum’ (Lso) is an unculturable phloem-limited bacterium that spreads from infected to healthy plants by psyllid vectors. This phytopathogenic bacterium includes strains responsible for diseases of several solanaceous and apiaceous crops (Haapalainen, 2014). Within this bacterial species, strains were classified into five haplotypes (LsoA, LsoB, LsoC, LsoD and LsoE). These haplotypes were defined based on genotyping of the 16S rRNA, 16S/23S ISR and 50S *rpIJ*-*rpIL* ribosomal gene loci, where any co-inherited single-nucleotide polymorphism (SNP) variant led to the classification in a new haplotype (Nelson et al., 2011, 2013; Teresani et al., 2014). LsoA and LsoB are associated with diseases of solanaceous crops, especially Zebra chip (ZC) disease of potato (*Solanum tuberosum*) in North America and New Zealand (Hansen et al., 2008; Liefting et al., 2008a; Lin el al., 2009), and are vectored to solanaceous plants by the potato/tomato psyllid, *Bactericera cockerelli* (Hemiptera: Triozidae) (Munyaneza et al., 2007; Secor et al., 2009). In addition to ZC disease, LsoA and LsoB cause diseases on other economically important solanaceous hosts, including tomato (*Solanum lycopersicum*), pepper (*Capsicum annuum*), eggplant (*Solanum melongena*), tomarillo (*Solanum betaceum*), tobacco (*Nicotiana tabacum)* and Cape gooseberry (*Physalis peruviana*) (Liefting et al., 2008b, Ling et al., 2011; Munyaneza et al., 2016). LsoC is vectored by the carrot psyllid *Trioza apicalis*, and is associated with carrot in northern Europe (Munyaneza et al., 2010a,b; 2012a,b; 2015; Nissinen et al., 2014). LsoD and LsoE are associated with the carrot psyllid *Bactericera trigonica*, and with vegetative disorders on several apiaceous crops in southwestern Europe and the Mediterranean area (Alfaro-Fernández et al., 2012a,b; Nelson et al., 2013; Loiseau et al., 2014; Tahzima et al., 2014, 2017; Teresani et al., 2014; Hajri et al., 2017; Mawassi et al., 2018). A best understanding of the relationships between Lso strains isolated from a set of various host plants in different geographical locations may help to better explain the increasing spread and extended host range of Lso.

Up to now, there has been no known biological mechanism explaining the different host range of the five haplotypes within Lso. The diversification of this species into different lineages could result from ecological adaptation, geographical structure or neutral processes (Haapalainen, 2014; Munyaneza, 2015). Genetic diversity investigations may provide an understanding of the population structure of a bacterial species and the evolutionary forces that led to its diversification (Puttamuk et al., 2014). For Lso, few studies have been attempted to investigate its population structure and genetic diversity, in particular due to its fastidious nature (Lin and Gudmestad, 2013). Sequencing of the 16S rRNA, 16S/23S ISR and 50S *rpIJ*-*rpIL* ribosomal gene loci has become the method of choice for the study of the genetic diversity of the bacterium (Lin et al., 2009; Wen et al., 2009; Nelson et al., 2011, 2013; Teresani et al., 2014; Alfaro-Fernández et al., 2017; Haapalainen et al., 2017; Hajri et al., 2017; Monger and Jeffries, 2018). However, genetic variation within these genes does not generally allow differentiation of closely related strains due to their high degree of conservation (Lin and Gudmestad, 2013). In addition to ribosomal genes, simple repeat sequences (SSR) and multilocus sequence typing (MLST) markers have been used to assess the genetic diversity of the bacterium (Lin et al., 2012; Glynn et al., 2012). However, these studies focused only on Lso solanaceous haplotypes (LsoA and LsoB), which is not sufficient to understand the extent of the genetic diversity of this bacterial species (Lin and Gudmestad, 2013; Haapalainen, 2014). More recently, genetic variation of LsoC was investigated by MLST (Haapalainen et al., 2018). To our knowledge, there is no information regarding the phylogenetic relationships between a large set of strains representing the five Lso haplotypes using multiple loci.

Multilocus sequence analysis (MLSA) has become the standard for phylogenetic analyses of bacterial species and has been shown to be a powerful molecular method for microbial population genetic studies. This method consists of the analysis of multiple (usually four to eight) conserved housekeeping genes, which encode proteins that are essential for the survival of the organism (Cooper and Feil, 2004; Gevers et al., 2005; Almeida et al., 2010). MLSA relies on the concatenation of aligned DNA sequences from each gene. Mutations within housekeeping genes are largely assumed to be selectively neutral, and therefore are more likely to correctly reflect the phylogeny of the strains (Gevers et al., 2005; Hajri et al., 2012). In addition, the concatenated sequence data could reduce the weight of horizontal gene transfer ‘HGT’ (Macheras et al., 2011) and/or recombination (Timilsina et al., 2015), which may provide more reliable phylogenetic relationships among closely related strains (Glaeser and Kämpfer, 2015). MLSA has been successfully used to describe the genetic structure of several phytopathogenic bacteria (Castillo and Greenberg, 2007; Young et al., 2008; Almeida et al., 2010; Trantas et al., 2013; Tancos et al., 2015; Constantin et al., 2016). Despite the availability of several Lso genomic sequences (Lin et al., 2011; Thompson et al., 2015; Wu et al., 2015; Wang et al., 2017), this strategy has never been applied to Lso.

In this paper, we studied the genetic structure of Lso and the phylogenetic relationships of strains belonging to different haplotypes that have so far been defined in this species. For this purpose, we designed a new MLSA scheme based on partial sequencing of seven housekeeping genes (*acnA, atpD, ftsZ, glnA, glyA, gnd* and *groEL*). Using a representative bacterial collection of 49 Lso strains, we evaluated its robustness with respect to defining the genetic structure of the bacterium.

## Materials and methods

### Bacterial strains

The 49 Lso strains used in this study are listed in Table 1. Strains were selected in order to maximize the diversity in terms of geographical origin, host and year of isolation. Forty-two strains have been isolated from several solanaceous and apiaceous crops and from different psyllids (*B. cockerelli, B. trigonica* and *T. apicalis*). In addition, we included for comparative purposes, the six published Lso genomes: strain ZC1 (Lin et al., 2011), strain RSTM (Wu et al., 2015), strain NZ1 and strain HenneA (Thompson et al., 2015), and strains FIN111 and FIN114 (Wang et al., 2017). We also included in our phylogenetic analysis the recently deposited draft genome sequence of LsoD (GenBank accession number NZ_PKRU00000000.1).

**Table 1.**
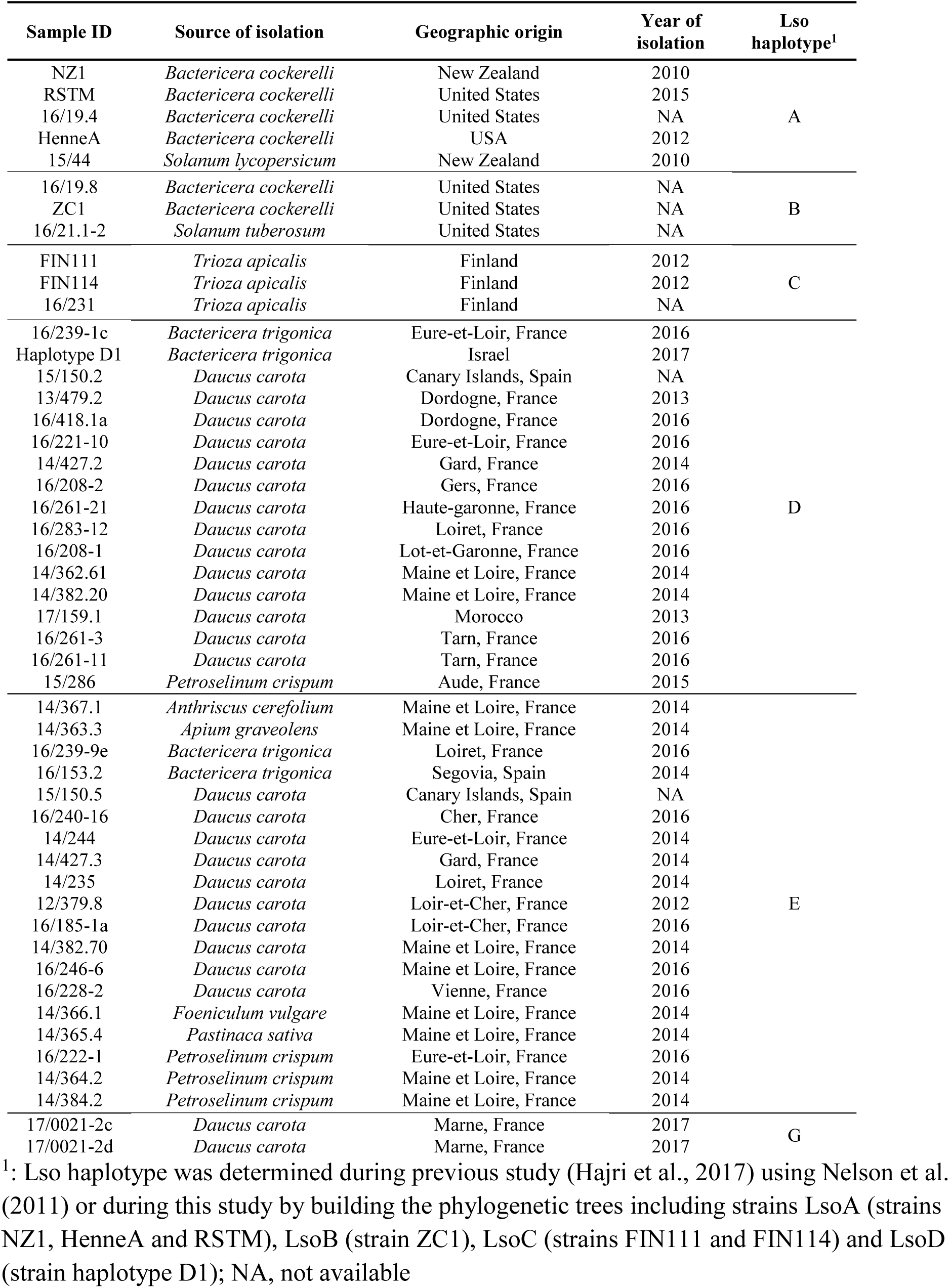
List of Lso strains used in this study

### DNA extraction

In this study, genomic DNAs of the 42 strains were extracted using (i) a slightly modified cetyltrimethylammonium bromide (CTAB) buffer extraction method (Murray and Thompson, 1980); and (ii) the magnetic-bead-based QuickPick™ SML Plant DNA Kit (Bio-Nobile) according to the manufacturer’s instructions. Each extraction series contained positive and negative controls.

### Selection of housekeeping genes

For the design of our MLSA scheme, several criteria were used in the selection of the potential genes. The genes included were (i) those encoding for putative housekeeping products with important biological functions (ii) evenly distributed across the genome as assessed from the Lso sequenced genomes, and (iii) present in one single copy in the Lso sequenced genomes. Using these criteria, we selected seven housekeeping genes: *acnA* (aconitate hydratase); *atpD* (ATP synthase subunit beta); *ftsZ* (cell division protein); *glnA* (glutamine synthetase I); *glyA* (serine hydroxymethyltransferase); *gnd* (6-phosphogluconate dehydrogenase) and *groEL* (molecular chaperone). Figure 1 shows the genomic location of the genes used in this study based on the Lso ZC1 genome.

**Figure 1.**
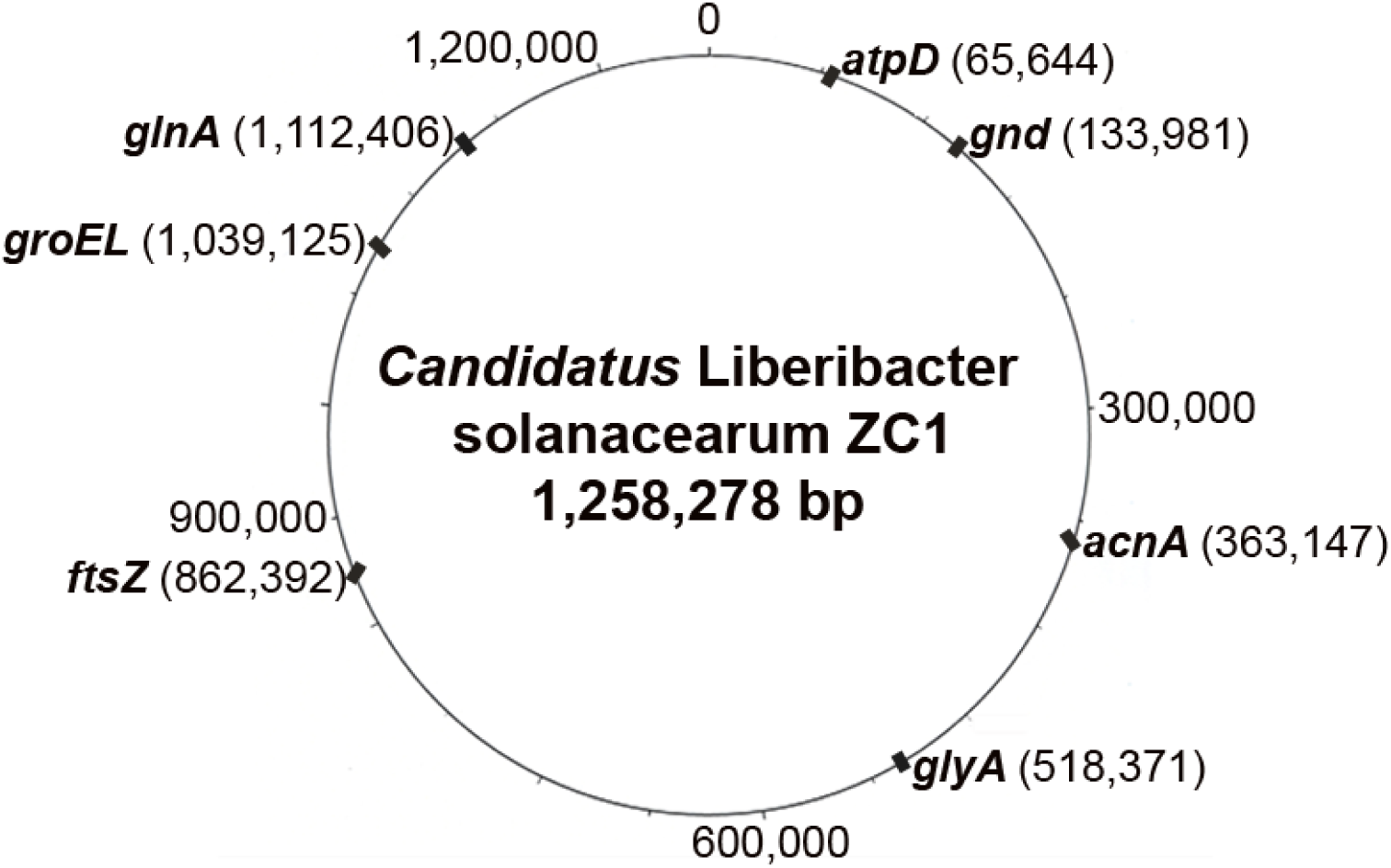
Schematic representation of the positions of the seven housekeeping genes used in 718 this study based on the sequenced Lso ZC1 genome (Genbank accession number CP002371). The position of each locus (in base pairs) is given between brackets.

### Gene amplification and sequencing

Primers for partial sequencing of the seven housekeeping genes were designed (Table 2) based on the alignments of orthologous sequences collected from the six Lso genomes available in GenBank: strain ZC1, strain NZ1, strain HenneA, strain RSTM, strain FIN111 and strain FIN114. PCR amplification of each gene was performed in a 25-µL reaction mix using the Bio-X-Act short polymerase mix (Bioline, London, UK), 0.2 μM of forward and reverse primers, and 2 μL of DNA. The PCR conditions were an initial denaturation at 95°C for 5 min; followed by 35 cycles of 30 s of denaturation at 94°C, 30 s of annealing at 58°C, and 1 min of extension at 72°C and followed by 10 min of a final extension at 72°C. All PCR series included positive and negative controls. Amplified PCR products were separated on 1.5% agarose gels containing ethidium bromide for visualization. PCR products were directly sequenced with forward and reverse primers (Genewiz, Takeley, UK).

**Table 2.**
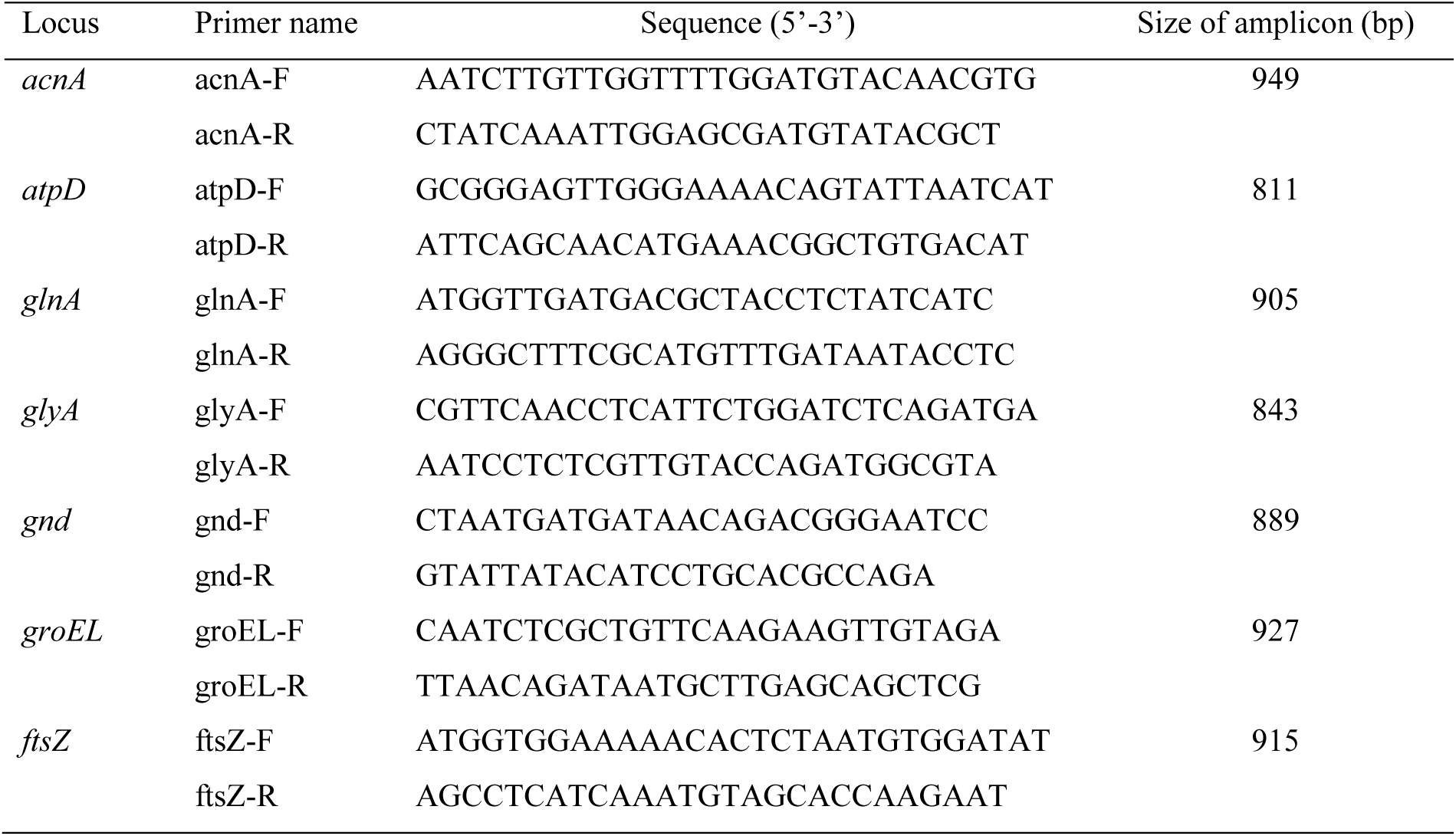
Primers of housekeeping genes used for amplification and sequencing

### Sequence acquisition and alignment

Forward and reverse nucleotide sequences were assembled using the CAP3 contig assembly program (Huang and Madan, 1999) to obtain high-quality sequences. Multiple alignments were edited using the ClustalW tool of the BIOEDIT program version 7.2.6.1 (Hall, 1999). Sequences were concatenated following the alphabetic order of the genes, ending in a sequence of 5415 bp (bp 1 to 816 for *acnA*, 817 to 1575 for *atpD*, 1576 to 2349 for *ftsZ*, 2350 to 3132 for *glnA*, 3133 to 3879 for *glyA*, 3880 to 4623 for *gnd*, and 4624 to 5415 for *groEL*).

### Sequence data analysis

Summary statistics for the sequences were computed using DnaSP version 5.10.01 (Librado and Rozas, 2009). The GC content, number of polymorphic sites (S), nucleotide diversities (θ*π* and θ*w*) (Nei, 1987; Watterson, 1975), and neutrality test of Tajima’s D (Tajima, 1989) were estimated for each of the seven genes and on concatenated sequences. DnaSP was also used to calculate the ratio of non-synonymous to synonymous substitutions (d*N*/d*S*) (Nei and Gojobori, 1986) in order to test the degree of selection on a locus. Values of d*N*/d*S* of 1, d*N*/d*S*>1 and d*N*/d*S* <1 indicate neutrality, diversifying selection and purifying selection, respectively. Occurrence of recombination was analyzed on each gene using the different recombination detection methods as implemented in RDP v4 (Martin et al., 2015). The analysis was performed with default settings for the different detection methods and the Bonferroni-corrected P-value cut-off was set at 0.05.

### Phylogenetic analyses

Phylogenetic analyses were performed on individual gene sequences as well as on the dataset of concatenated sequences of the 49 strains of our bacterial collection. Strain psy62 of ‘*Ca.* L. asiaticus’ (Duan et al., 2009) was used to root trees. Maximum Likelihood trees were generated with the MEGA 7.0.14 software program (Kumar et al., 2016) using the Kimura two-parameter model (Kimura, 1980). Statistical support for tree nodes was evaluated by bootstrap analyses with 1000 replicates. To determine the haplotype affiliation of tested Lso strains, the sequenced genomes of strains from LsoA (strains NZ1, HenneA and RSTM), LsoB (strain ZC1), LsoC (strains FIN111 and FIN114) and LsoD (strain haplotype D1) were included in the analysis. For LsoE for which no published genome sequence is available, we used as reference, strain 14/235 that was previously determined as belonging to LsoE using the 16S rRNA gene and the 50S ribosomal protein *rpIJ-rpIL* gene region (Hajri et al., 2017).

### Sequence accession numbers

Two hundred and ninety four sequences generated in this study were deposited in the GenBank database with accession numbers MH108648 - MH108689 for *acnA*, MH108690 - MH108731 for *atpD*, MH108732 - MH108773 for *ftsZ*, MH108774 - MH108815 for *glnA*, MH108816 - MH108857 for *glyA*, MH108858 - MH108899 for *gnd* and MH108900 - MH108941 for *groEL.*

## Results

### Sequence analysis and DNA polymorphism

In the present study, amplification was successful for all tested Lso strains with the seven housekeeping genes. To validate the choice of the seven loci as appropriate phylogenetic markers for Lso, descriptive statistics on nucleotide and allelic diversities were calculated for each locus and for the concatenated dataset. The 49 sequences of each locus were aligned; no gap and no insertion in the sequences of all tested Lso strains were detected. Analyzed sequence lengths ranged from 744 bp (*gnd*) to 816 bp (*acnA*), leading to a total of 5415 bp for the sequence of concatenated dataset (Table 3). The GC content for all loci ranged from 33.9% (*gnd*) to 43.2% (*ftsz*). All loci were polymorphic and the number of polymorphic nucleotide sites varied from 12 for the least polymorphic locus (*ftsZ*) to 27 for the most polymorphic locus (*glnA*) (Table 3). The value for the neutrality test of Tajima was positive for 4 loci (*ftsZ, glnA, glyA* and *groEL*), negative for 3 loci (*acnA, atpD* and *gnd*) and positive for the concatenated data but it was not significant for all tested loci and for concatenated data. The d*N*/d*S* ratios of all tested housekeeping genes were <1, indicating that these loci were subject to purifying selection. No recombination event was detected with the RDP program, indicating that the Lso population is highly clonal. For the concatenated dataset, the average GC content was 38.9%. Overall, 143/5415 nucleotide sites examined here are polymorphic, which indicates that Lso is genetically rather uniform. For neutrality tests, no significant departure from neutrality was observed on the overall dataset with the Tajima’s D test (Table 3).

**Table 3.**
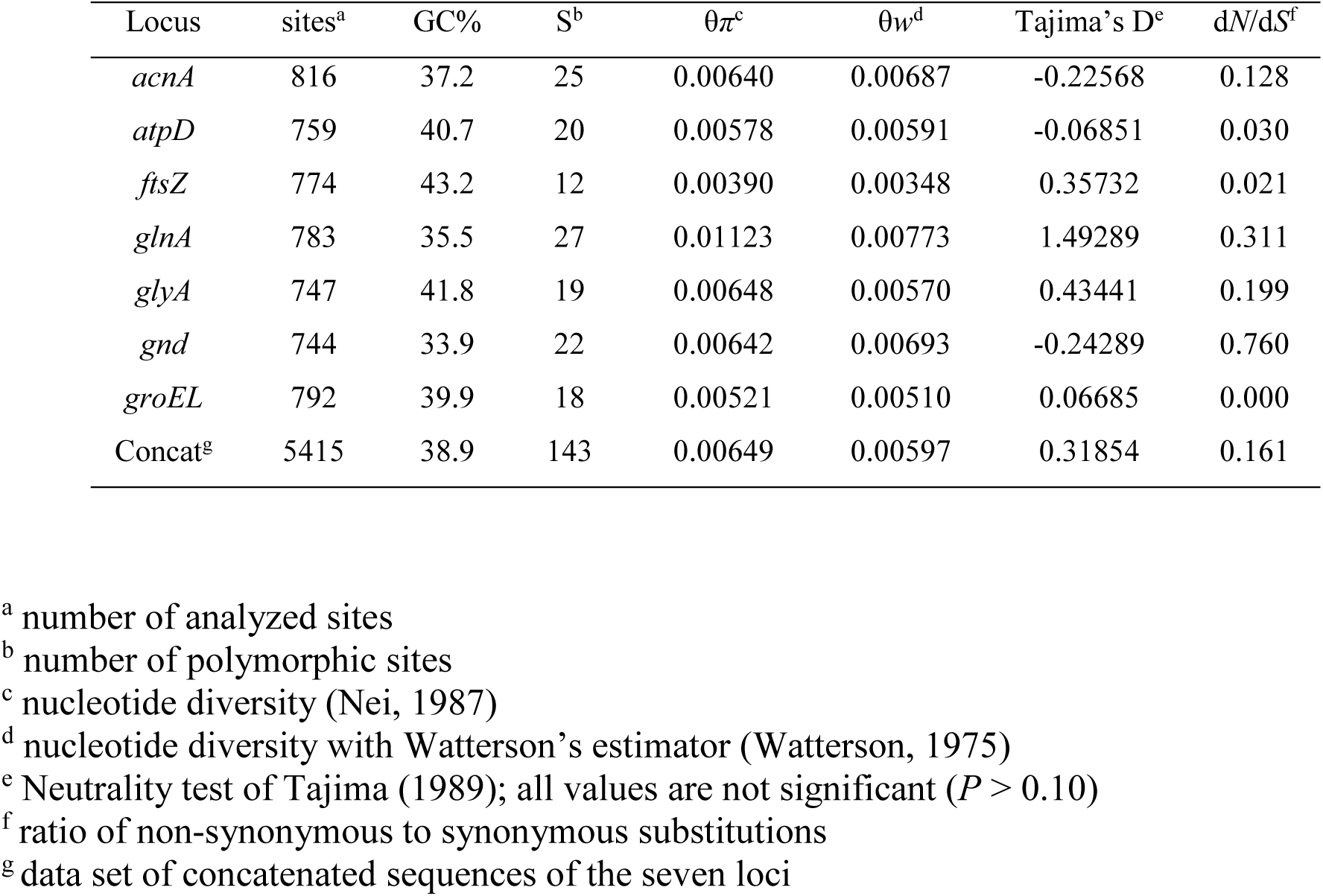
Summary statistics for the seven housekeeping genes and concatenated sequences used in this study

### MLSA based on concatenated gene sequences

The phylogenetic tree generated from the concatenated sequences of the seven housekeeping genes (*acnA, atpD, ftsZ, glnA, glyA, gnd* and *groEL*) showed six well defined groups. High bootstrap values indicated that this clustering was well supported and that the phylogenetic tree was robust (Figure 2). The first group (LsoB) is composed of three strains isolated in the United States: one from *S. tuberosum*, one from *B. cockerelli* and the sequenced strain ZC1. The second group (LsoC) consists of one strain isolated in Finland from *T. apicalis* and the sequenced strains FIN111 and FIN114. The third group (LsoA) was identified from one strain isolated from *B. cockerelli* in the United States, one strain isolated from *S. lycopersicum* in New Zealand, and the sequenced strains NZ1, RSTM and HenneA. The fourth group (LsoD) included strain haplotype D1 from Israel, one strain isolated in Spain from carrot plants, one strain isolated in Morocco from carrot plants, and 14 strains isolated in France: 12 from carrot plants, one from *B. trigonica* and one from parsley. The fifth group (LsoE) included two strains isolated from Spain: one from *B. trigonica* and one from carrot plants, and 17 strains isolated from France (nine from carrot plants, one from *B. trigonica*, three from parsley, one from celery, one from fennel, one from parsnip and one from chervil) (Table 1). In addition to these five groups, which correspond to the five Lso haplotypes described so far, our phylogenetic tree revealed the presence of a sixth haplotype, designated LsoG. This new haplotype clustered two strains from France (17/0021-2c and 17/0021-2d). The bootstrap value (95) of the LsoG group depicted the robustness of this phylogenetically novel population (Figure 2).

**Figure 2.**
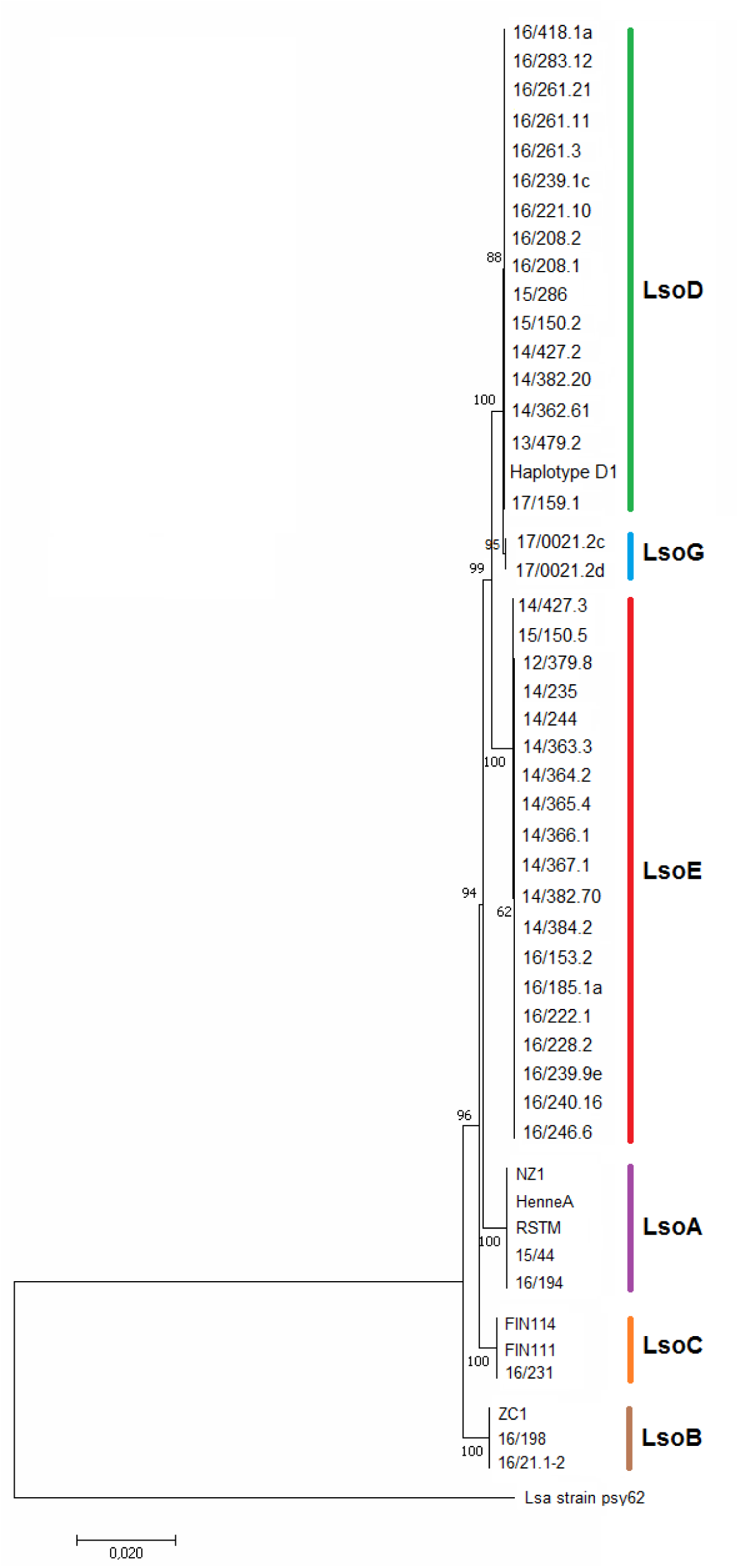
Phylogenetic tree of the MLSA based on concatenated genes (*acnA, atpD, ftsZ, glnA, glyA, gnd* and *groEL*) among the 49 Lso strains. The tree is based on 5415 bp of common sequence. The tree was constructed using the maximum likelihood method. The confidence of the nodes was tested with 1,000 bootstrap replicates. The scale bar indicates the number of nucleotide substitutions per site. The tree is rooted with the ‘*Ca.* L. asiaticus’ strain psy62.

Regarding the phylogenetic relationships between the 49 tested Lso strains, a close relatedness was observed between LsoD, LsoE and LsoG. These three haplotypes appeared to be more closely related to LsoA than to LsoC. Strains of LsoB formed a cluster that was the most genetically distant from the other five Lso haplotypes (Figure 2). Based on our MLSA analysis, strains that are pathogenic on a same plant species or family were displayed in very divergent groups. This was the case of LsoA and LsoB, both pathogenic on solanaceous crops, which did not group together on the basis of our sequence data. For apiaceous haplotypes, LsoC form a distinct phylogenetic group that was clearly separated from the three other apiaceous haplotypes (LsoD, LsoE and LsoG) (Figure 2). These observations suggest that host specificity is not correlated to Lso classification based on our phylogenetic data.

### Individual gene phylogenies

Individual phylogenetic trees were constructed using the maximum likelihood method and are shown in Figures S1 to S7. The tree topologies constructed from single genes presented roughly congruent phylogenies. Some differences were observed in the topology of the individual phylogenetic trees. For example, LsoC appears to be more closely related to the other apiaceous haplotypes in the phylogenetic trees built with the *acnA, glnA* and *glyA* genes. This was not the case for the trees built with the four other genes (*atpD, ftsZ, gnd* and *groEL*) since LsoC form a cluster that was phylogenetically distant from the cluster formed by the other apiaceous haplotypes.

All the individual phylogenetic trees confirmed the structuration of the tested Lso strains into five groups (LsoA, LsoB, LsoC, LsoD and LsoE), with the formation of a sixth group (LsoG) in the *glnA, glyA, groEL* and *ftsZ* phylogenetic trees (Figures S1 to S7). The gene sequences from LsoG strains had distinct sequences in these four housekeeping genes: they exhibited one new SNP in the *glnA* gene, one new SNP in the *glyA* gene and one new SNP in the *groEL* gene (Table 4). In addition, the *ftsZ* gene sequences from these two strains varied from the *ftsZ* gene sequences of the other LsoD strains by two nucleotide substitutions (Table 4). Regarding LsoE, all tested strains have identical sequences except for the *atpD* gene (Figures S1 to S7), for which two strains (14/427.3 from France and 15/150.5 from Spain) exhibited a different allele from the other LsoE strains (presence of G instead of T at position 321 of the aligned sequence of the *atpD* gene).

**Table 4.**
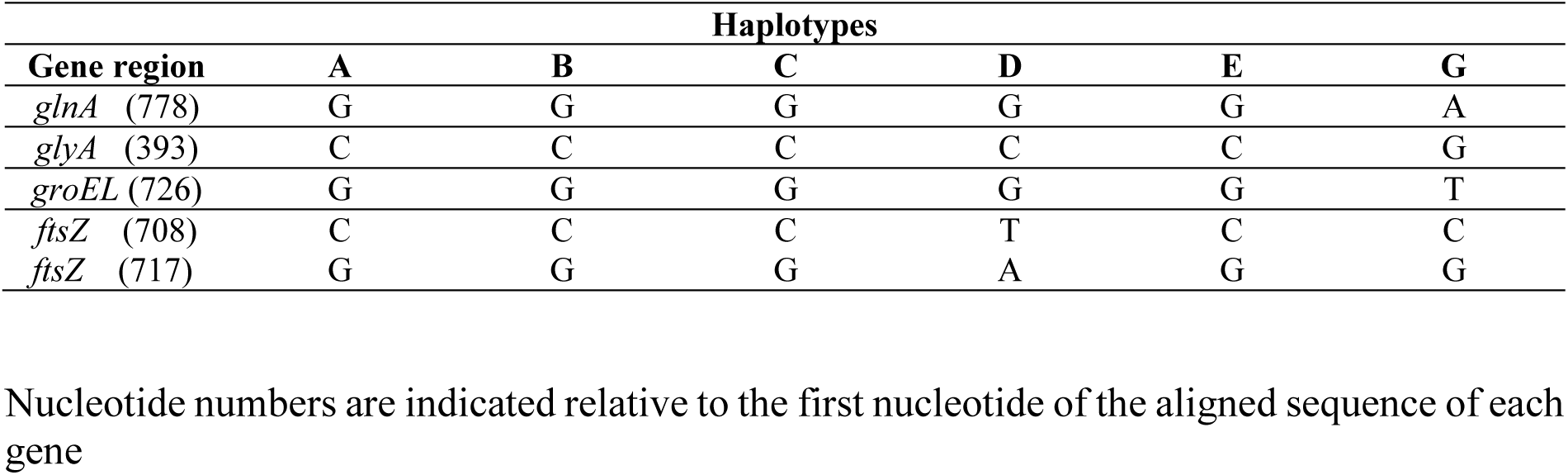
New SNP features of the LsoG haplotype

## Discussion

Lso is an emerging phytopathogenic bacterium which is responsible for several economically important diseases on solanaceous and apiaceous crops worldwide (Soliman et al., 2013; Haapalainen, 2014). Previous sequence-based analyses of the genetic diversity of Lso have only targeted genes of the ribosomal operon (Lin et al., 2009; Wen et al., 2009; Nelson et al., 2011). These data, however, were obtained based on the analysis of a single locus, which may not provide enough information for discrimination between different Lso strains (Lin and Gudmestad, 2013). Since the publication of the first Lso genome sequence (Lin et al., 2011), a number of loci including SSR and MLST markers were used to define the genetic relationships among solanaceous Lso haplotypes (LsoA and LsoB) (Lin et al., 2012; Glynn et al., 2012). A more recent study investigated the genetic variability of LsoC strains in Finland by MLST (Haapalainen et al., 2018). These genetic diversity studies did not take into account LsoD and LsoE variability. Thus, an updated phylogeny of Lso strains representing the five established haplotypes is lacking. In recent years, the MLSA approach has shed new light on prokaryotic phylogeny, and has been widely used in the classification and identification of diverse bacterial species (Gevers et al., 2005; Almeida et al., 2010; Glaeser and Kämpfer, 2015). In this study, we established a new MLSA scheme based on seven housekeeping genes (*acnA, atpD, ftsZ, glnA, glyA, gnd* and *groEL*) for a representative collection of 49 Lso strains. The availability of 5415 bp of sequence data from each of the 49 strains gives an important resource to understand the extent and nature of genetic diversity within this bacterial species.

Considering the seven housekeeping genes used in this study, 143 polymorphic sites in 5415 bp of sequence were identified. No insertions/deletions were found in the analyzed sequences. The seven loci were informative, with *glnA* being the most variable gene. In addition to point mutations, bacterial evolution may be driven by recombination which is considered as one of the most important processes that increase genetic variability in bacterial genomes (Feil and Spratt, 2001; Martin et al., 2011). Due to the association of LsoA and LsoB with the same solanaceous crops and the same vector (*B. cockerelli*) (Munyaneza et al., 2007; Lin et al., 2012), recombination between these two haplotypes could occur. Within apiaceous haplotypes, we would expect recombination to occur between LsoD and LsoE since they are both associated with *B. trigonica* (Alfaro-Fernández et al., 2012b; Teresani et al., 2014; Tahzima et al., 2017) whereas LsoC is vectored by a different psyllid species (*T. apicalis*) (Munyaneza et al., 2010b). Recombination events were estimated on each locus. No recombination events were detected whatever the data analyzed. The absence of recombination for housekeeping genes was previously reported for other phytopathogenic bacteria such as *Ralstonia solanacearum* and *Xanthomonas* species (Castillo and Greenberg, 2007; Bui Thi Ngoc et al., 2010; Hajri et al., 2012). Housekeeping genes are components of the ‘core genome’ and typically encode proteins essential for the organism’s survival. These genes are present in all strains of a bacterial species and usually evolve slowly (Sarkar and Guttman, 2004). In the frame of this study, the levels of selection acting on the tested housekeeping genes were evaluated with two population genetic tests. Tajima’s D statistic did not produce significant results. In addition, all d*N*/d*S* were less than 1, indicating that the seven tested loci appeared to be under purifying selection, as expected for genes chosen for MLSA studies. Taken together, our data revealed a high clonality for housekeeping genes and support a predominant role of mutation in shaping the genetic diversity of Lso.

Previous genetic studies based on the sequencing of three ribosomal gene loci (16S rRNA, 16S/23S ISR and 50S *rpIJ*-*rpIL*) led to consider Lso as a species complex constituted of five haplotypes (LsoA, LsoB, LsoC, LsoD and LsoE) (Nelson et al., 2011, 2013; Teresani et al., 2014). Our MLSA data showed that differences between the five haplotypes are not restricted to the rRNA operon and are present within the core genome. The phylogenetic tree based on concatenated sequences of the seven housekeeping genes confirmed the clustering of Lso strains into five evolutionary lineages that correspond to the five previously described Lso haplotypes, supporting the hypothesis of Nelson et al (2011) that Lso haplotypes represent stable haplotypes. This clear correspondence between phylogenetic clustering and haplotype classification was also supported by phylogenetic trees based on individual loci. The *atpD, ftsZ, gnd* and *groEL* phylogenetic trees were found to be congruent with the tree generated from the concatenated data, indicating a common evolutionary history for these four genes. However, the *acnA, glnA* and *glyA* gene phylogenies did not accurately represent the same evolutionary history as the other tested genes. The main incongruence concerned the position of LsoA, LsoB and LsoC. A recent phylogenetic analysis based on 88 single-copy orthologs of the bacterial core genome showed that LsoA is more closely related to LsoC than to LsoB (Wang et al., 2017). This topology is in agreement with the *atpD, ftsZ, gnd* and *groEL* phylogenies, indicating that these genes are more reliable to reflect the phylogenetic relationships among Lso haplotypes. In addition, our phylogenetic data revealed a clear separation between the cluster formed by LsoD and LsoE on the one hand and the cluster formed by LsoC on the other hand, although displaying the same host range. Similarly, LsoA and LsoB which are both associated with solanaceous crops, did not cluster together on the basis of our phylogenetic analysis. Thus, no clear correspondence could be established between clustering of Lso strains and their host specificities on the basis of our phylogenetic data. When considering the insect vector, our phylogenetic analysis suggests that clustering of Lso strains associated with apiaceous crops may be explained by differences in insect vector. Indeed, LsoD and LsoE, which appeared to be very close phylogenetically, share the same insect vector (*B. trigonica*) (Alfaro-Fernández et al., 2012b; Teresani et al., 2014; Tahzima et al., 2017), whereas LsoC which forms a more distant cluster, is associated with a different psyllid species (*T. apicalis*) (Munyaneza et al., 2010b). The genetic relatedness of LsoD and LsoE was reported previously based on sequence analysis of the 50S *rpIJ*-*rpIL* gene (Hajri et al., 2017). For solanaceous haplotypes, this is not true since LsoA and LsoB, which are both associated with *B. cockerelli*, did not cluster together on the basis of our phylogenetic analysis. Altogether, the results presented here strongly suggest that the current host plant range and the vector psyllid associations of the different Lso haplotypes are not congruent with Lso phylogeny, as previously suggested by Haapalainen (2014).

The present study not only clarifies the phylogenetic relationships among Lso haplotypes but also reveals a part of the unknown genetic diversity of the bacterium. In addition to the five previously described Lso haplotypes (LsoA, LsoB, LsoC, LsoD and LsoE), our phylogenetic data revealed the presence of a sixth haplotype, designated LsoG, This new genetically distinct group is represented by two strains (17/0021-2c and 17/0021-2d) from France and is associated with asymptomatic carrots. Our sequence analysis revealed that the new haplotype G had distinct sequences in four out of the seven tested housekeeping genes: one new SNP in the *glnA* gene, one new SNP in the *groEL* gene, one new SNP in the *glyA* gene and one different allelic form in the *ftsZ* gene. In the phylogenetic tree based on the concatenated gene sequences, strains 17/0021-2c and 170021-2d form a very tight and homogeneous cluster which is clearly separate from the other Lso haplotypes. Because strains 17/0021-2c and 17/0021-2d were isolated from the same geographical region, it is tempting to speculate that the newly described haplotype G is a genetically structured population based on geographical area. To support this hypothesis, further characterization of additional representative strains of LsoG is now necessary. Although the haplotype nomenclature system has been useful in an agricultural context, its biological justification is questionable since there are no biological distinctions that reliably differentiate Lso haplotypes (Lin and Gudmestad, 2013). Thus, haplotype designation within Lso should take into account both the genetic diversity and biological traits of the strains (Haapalainen, 2014; Munyaneza, 2015). More recently, two novel haplotypes (LsoU and LsoF) was identified in Finland from the psyllid *Trioza urticae* and its host plant *Urtica dioica*, and in the USA in potato tuber (Haapalainen et al., 2018; Swisher-Grimm and Garczynski, 2018). The identification of three novel haplotypes (LsoU in Finland, LsoF in USA and LsoG in France) indicates that genetic diversity of Lso is much higher than previously expected. In our study, we focused on housekeeping genes which provide substantial evidence that strains 17/0021-2c and 17/0021-2d represent a new Lso haplotype. However, we cannot exclude the possibility that genetic variation among these strains is present elsewhere in the genome. Further work on the biological traits (*i.e.* host range) of these two strains and further sequencing of more variable genes like those involved in virulence are now necessary to confirm the status of this new haplotype.

The interaction between Lso and its host plants is strongly influenced by the ecological niche, the insect vector and the evolution of bacterial pathogenicity determinants (Lin and Gudmestad, 2013; Haapalainen, 2014). Currently, little is known about pathogenicity determinants that can account for the differing host specificities between Lso haplotypes. Previous comparative genomic studies focused only on LsoA, LsoB and LsoC and revealed relevant differences in the genome organization between these haplotypes (Lin et al., 2011; Thompson et al., 2015; Wang et al., 2017). However, these studies did not take into account haplotypes D and E since no published genome sequence is available for LsoE and a draft genome sequence of LsoD was made available only recently in Genbank. Notably, our MLSA data highlighted interesting features for LsoD and LsoE: (i) they are very closely related, but phylogenetically distinct; (ii) they form a cluster that is clearly separated from LsoC which is also pathogenic on apiaceous crops. In addition, a recent study reveals that the newly described LsoU haplotype is more closely related to LsoD and LsoA than to LsoC (Haapalainen et al., 2018). Altogether, these observations raise interesting questions about the genomic basis of adaptation to the host of Lso haplotypes. Sequencing of LsoE, LsoF, LsoU and LsoG genomes is now necessary to perform a whole comparative genomic analysis at the species level which may enable the identification of specific-haplotype genes. Such traits are excellent candidates to better understand the genetic basis of the host range differences that exist between apiaceous and solanaceous haplotypes and the factors involved in vector/plant host interactions. In this article, we have described the genetic structure of Lso with special emphasis on housekeeping genes. However, the population structure of a bacterium can also be tracked by focusing on other informative genetic markers such as phage-related genes that could be linked to virulence. The potential contribution of prophages and phage-related sequences to the genetic diversity of Liberibacter species has been investigated more extensively in ‘*Ca.* L. asiaticus’ (Zhou et al., 2013; Puttamuk et al., 2014; Zheng et al., 2016, 2018). Furthermore, Thompson *et al.*(2015) highlighted that the most important difference between the two genome sequences of LsoA is the location of their prophage domains. Profiling the phage-related genes of Lso strains from different geographic origin could help to better decipher the evolutionary forces responsible for the haplotype diversification within this bacterial species and to identify the sources of disease outbreaks or incursions.

## Author contributions

AH: conceived and designed the experiments, analyzed and interpreted the data, wrote the paper; PCS: performed the experiments; PG: revised the manuscript; ML: analyzed the data with the maximum likelihood method and revised the manuscript.

## Funding

This study was partially supported by funding from the European Union’s Horizon 2020 research and innovation programme under grant agreement No 635646: POnTE (Pest Organisms Threatening Europe) and by funding from the French ministry of agriculture under grant agreement C-2015-09 n°2101755701: CaLiso.

### Acknowledgments

We gratefully thank the partners of the CaLiso project (FNAMS, CTIFL, Fredon Centre, chambre d’Agriculture de la Marne, UFS) for providing us a part of the French Lso isolates used in this study. We also thank Joseph Munyaneza, Sarah Thompson, Anne Nissinen, Alberto Fereres, Felipe Siverio de la Rosa and Stephanie Mallard for kindly providing us with Lso strains from USA, New Zealand, Finland, Spain, Canary Islands and Morocco.

## Supplementary material

**Figure S1.** Phylogenetic tree based on the *acnA* gene among the 49 Lso strains. The tree was constructed using the maximum likelihood method. Bootstrap values over 50 (1,000 replicates) were shown at each node. The scale bar indicates the number of nucleotide substitutions per site. The tree is rooted with the ‘*Ca.* L. asiaticus’ strain psy62.

**Figure S2.** Phylogenetic tree based on the *atpD* gene among the 49 Lso strains. The tree was constructed using the maximum likelihood method. Bootstrap values over 50 (1,000 replicates) were shown at each node. The scale bar indicates the number of nucleotide substitutions per site. The tree is rooted with the ‘*Ca.* L. asiaticus’ strain psy62.

**Figure S3.** Phylogenetic tree based on the *ftsZ* gene among the 49 Lso strains. The tree was constructed using the maximum likelihood method. Bootstrap values over 50 (1,000 replicates) were shown at each node. The scale bar indicates the number of nucleotide substitutions per site. The tree is rooted with the ‘*Ca.* L. asiaticus’ strain psy62.

**Figure S4.** Phylogenetic tree based on the *glnA* gene among the 49 Lso strains. The tree was constructed using the maximum likelihood method. Bootstrap values over 50 (1,000 replicates) were shown at each node. The scale bar indicates the number of nucleotide substitutions per site. The tree is rooted with the ‘*Ca.* L. asiaticus’ strain psy62.

**Figure S5.** Phylogenetic tree based on the *glyA* gene among the 49 Lso strains. The tree was constructed using the maximum likelihood method. Bootstrap values over 50 (1,000 replicates) were shown at each node. The scale bar indicates the number of nucleotide substitutions per site. The tree is rooted with the ‘*Ca.* L. asiaticus’ strain psy62.

**Figure S6.** Phylogenetic tree based on the *gnd* gene among the 49 Lso strains. The tree was constructed using the maximum likelihood method. Bootstrap values over 50 (1,000 replicates) were shown at each node. The scale bar indicates the number of nucleotide substitutions per site. The tree is rooted with the ‘*Ca.* L. asiaticus’ strain psy62.

**Figure S7.** Phylogenetic tree based on the *groEL* gene among the 49 Lso strains. The tree was constructed using the maximum likelihood method. Bootstrap values over 50 (1,000 replicates) were shown at each node. The scale bar indicates the number of nucleotide substitutions per site. The tree is rooted with the ‘*Ca.* L. asiaticus’ strain psy62.

## References

Alfaro-Fernández, A., Cebrián, M. C., Villaescusa, F. J., Hermoso de Mendoza, A., Ferrándiz, J. C., Sanjuán, S., et al. (2012a). First report of ‘*Candidatus* Liberibacter solanacearum’ in carrots in mainland Spain. Plant Dis. 96:582. doi:10.1094/PDIS-11-11-0918-PDN

Alfaro-Fernández, A., Siverio, F., Cebrián, M. C., Villaescusa, F. J., and Font, M. I. (2012b). ‘*Candidatus* Liberibacter solanacearum’ associated with *Bactericera trigonica*-affected carrots in the Canary Islands. Plant Dis. 96:581. doi:10.1094/PDIS-10-11-0878-PDN

Alfaro-Fernández, A., Hernández-Llopis, D., and Font, M. I. (2017). Haplotypes of ‘*Candidatus* Liberibacter solanacearum’ identified in Umbeliferous crops in Spain. Eur. J. Plant Pathol. 149, 127–131. doi:10.1007/s10658-017-1172-2

Almeida, N. F., Yan, S., Cai, R., Clarke, C. R., Morris, C. E., Schaad, N. W., et al. (2010). PAMDB, a multilocus sequence typing and analysis database and website for plant-associated microbes. Phytopathology 100, 208-215. doi:10.1094/PHYTO-100-3-0208

Bui Thi Ngoc, L., Vernière, C., Jarne, P., Brisse, S., Guérin, F., Boutry, S., et al. (2009). From local surveys to global surveillance: three high throughput genotyping methods for the epidemiological monitoring of *Xanthomonas citri* pv. citri pathotypes. Appl. Environ. Microbiol. 75, 1173–1184. doi:10.1128/AEM.02245-08

Castillo, J. A., and Greenberg, J. T. (2007). Evolutionary dynamics of *Ralstonia solanacearum*. Appl. Environ. Microbiol. 73, 1225–1238. doi:10.1128/AEM.01253-06

Constantin, E. C., Cleenwerck, I., Maes, M., Baeyen, S., Van Malderghem, C., De Vos, P., et al. (2016). Genetic characterization of strains named as *Xanthomonas axonopodis* pv. dieffenbachiae leads to a taxonomic revision of the X. axonopodis species complex. Plant Pathol. 65, 792–806. doi:10.1111/ppa.12461

Cooper, J. E., and Feil, E. J. (2004). Multilocus sequence typing - what is resolved. Trends Microbiol. 12, 373-377. doi:10.1016/j.tim.2004.06.003

Duan, Y. P., Zhou, L. J., Hall, D. G., Li, W. B., Doddapaneni, H., Lin, H., et al. (2009). Complete genome sequence of citrus huanglongbing bacterium, ‘*Candidatus* Liberibacter asiaticus’ obtained through metagenomics. Mol. Plant Microbe Interact. 22, 1011–1020. doi:10.1094/MPMI-22-8-1011

Feil, E. J., and Spratt, B. G. (2001). Recombination and the population structures of bacterial pathogens. Annu. Rev. Microbiol. 55, 561–590. doi:10.1146/annurev.micro.55.1.561

Gevers, D., Cohan, F. M., Lawrence, J. G., Spratt, B. G., Coenye, T., Feil, E. J., et al. (2005). Re-evaluating prokaryotic species. Nat.Rev. Microbiol. 3, 733–739. doi:10.1038/nrmicro1236

Glaeser, S.P., and Kämpfer, P. (2015). Multilocus sequence analysis (MLSA) in prokaryotic taxonomy. Syst. Appl. Microbiol. 38, 237–245. doi:10.1016/j.syapm.2015.03.007

Glynn, J. M., Islam, M. S., Bai, Y., Lan, S., Wen, A., Gudmestad, N. C., et al. (2012). Multilocus sequence typing of ‘*Candidatus* Liberibacter solanacearum’ isolates from North America and New Zealand. J. Plant Pathol. 94, 223–228. doi:10.4454/jpp.fa.2012.007

Haapalainen, M. (2014). Biology and epidemics of ‘*Candidatus* Liberibacter’ species, psyllid-transmitted plant-pathogenic bacteria. Ann. Appl. Biol. 165, 172–198. doi:10.1111/aab.12149

Haapalainen, M., Kivimäki, P., Latvala, S., Rastas, M., Hannukkala, A., Jauhiainen, L., et al. (2017). Frequency and occurrence of the carrot pathogen ‘*Candidatus* Liberibacter solanacearum’ haplotype C in Finland. Plant Pathol. 66, 559–570. doi:10.1111/ppa.12613

Haapalainen, M., Wang, J., Latvala, S., Lehtonen, M. T., Pirhonen, M., and Nissinen, A. I. (2018). Genetic variation of ‘*Candidatus* Liberibacter solanacearum’ haplotype C and identification of a novel haplotype from *Trioza urticae* and stinging nettle. Phytopathology, “in press”. doi:10.1094/PHYTO-12-17-0410-R

Hajri, A., Brin, C., Zhao, S., David, P., Feng, J.-X., Koebnik, R., et al. (2012). Multilocus sequence analysis and type III effector repertoire mining provide new insights into the evolutionary history and virulence of *Xanthomonas oryzae*. Mol. Plant Pathol. 13, 288–302. doi: 10.1111/j.1364-3703.2011.00745.x

Hajri, A., Loiseau, M., Cousseau-Suhard, P., Renaudin, I., and Gentit, P. (2017). Genetic characterization of ‘*Candidatus* Liberibacter solanacearum’ haplotypes associated with apiaceous crops in France. Plant Dis. 101, 1383–1390. doi:10.1094/PDIS-11-16-1686-RE

Hall, T. A. (1999). BioEdit: a user-friendly biological sequence alignment editor and analysis program for Windows 95/98/NT. Nucleic Acids Symp. Ser. 41, 95–98.

Hansen, A. K., Trumble, J. T., Stouthamer, R., and Paine, T. D. (2008). A new Huanglongbing (HLB) species, “*Candidatus* Liberibacter psyllaurous”, found to infect tomato and potato, is vectored by the psyllid *Bactericera cockerelli* (Sulc). Appl. Environ. Microbiol. 74, 5862–5865. doi:10.1128/AEM.01268-08

Huang, X., and Madan, A. (1999). CAP3: A DNA sequence assembly program. Genome Res. 9, 868–877. doi:10.1101/gr.9.9.868

Kimura, M. (1980). A simple method for estimating evolutionary rates of base substitutions through comparative studies of nucleotide sequences. J. Mol. Evol. 16, 111–120.

Kumar, S., Stecher, G., and Tamura, K. (2016). MEGA7: Molecular Evolutionary Genetics Analysis version 7.0 for bigger datasets. Mol. Biol. Evol. 33: 1870–1874. doi:10.1093/molbev/msw054

Librado, P., and Rozas, J. (2009). DnaSP v5: a software for comprehensive analysis of DNA polymorphism data. Bioinformatics 25, 1451–1452. doi:10.1093/bioinformatics/btp187

Liefting, L., Perez-Egusquiza, Z., Clover, G., and Anderson, J. (2008a). A new ‘*Candidatus* Liberibacter’ species In Solanum tuberosum in New Zealand. Plant Dis. 92:1474. doi:10.1094/PDIS-92-10-1474A

Liefting, L. W., Ward, L. I., Shiller, J. B., and Clover, G. R. G. (2008b). A new ‘*Candidatus* Liberibacter’ species In Solanum betaceum (tamarillo) and *Physalis peruviana* (cape gooseberry) in New Zealand. Plant Dis. 92:1588. doi:10.1094/PDIS-92-11-1588B

Lin, H., and Gudmestad, N. C. (2013). Aspects of Pathogen genomics, diversity, epidemiology, vector dynamics, and disease management for a newly emerged disease of potato: zebra chip. Phytopathology 103, 524–537. doi:10.1094/PHYTO-09-12-0238-RVW

Lin, H., Doddapaneni, H., Munyaneza, J. E., Civerolo, E. L., Sengoda, V. G., Buchman, J. L., et al. (2009). Molecular characterization and phylogenetic analysis of 16S rRNA from a new “*Candidatus* Liberibacter” strain associated with zebra chip disease of potato (*Solanum tuberosum* L.) and the potato psyllid (*Bactericera cockerelli* Sulc). J. Plant Pathol. 91, 215–219. doi:10.4454/jpp.v91i1.646

Lin, H., Lou, B., Glynn, J. M., Doddapaneni, H., Civerolo, E. L., Chen, C., et al. (2011). The complete genome sequence of ‘*Candidatus* Liberibacter solanacearum’, the bacterium associated with potato zebra chip disease. PLoS ONE 6:e19135. doi:10.1371/journal.pone.0019135

Lin, H., Islam, M. S., Bai, Y., Wen, A., Lan, S., Gudmestad, N. C., et al. (2012). Genetic diversity of ‘*Candidatus* Liberibacter solanacearum’ strains in the United States and Mexico revealed by simple sequence repeat markers. Eur. J. Plant Pathol. 132, 297–308. doi:10.1007/s10658-011-9874-3

Ling, K. S., Lin, H., Ivey, M. L. L., Zhang, W., and Miller, S. (2011). First report of ‘*Candidatus* Liberibacter solanacearum’ naturally infecting tomatoes in the state of Mexico, Mexico. Plant Dis. 95:1026. doi:10.1094/PDIS-05-11-0365

Loiseau, M., Garnier, S., Boirin, V., Merieau, M., Leguay, A., Renaudin, I., et al. (2014). First Report of ‘*Candidatus* Liberibacter solanacearum’ in carrot in France. Plant Dis. 98:839. doi:10.1094/PDIS-08-13-0900-PDN

Macheras, E., Roux, A. L., Bastian, S., Leao, S. C., Palaci, M., Sivadon-Tardy, V., et al. (2011). Multilocus sequence analysis and *rpoB* sequencing of *Mycobacterium abscessus* (Sensu Lato) strains. J. Clin. Microbiol. 49, 491–499. doi:10.1128/JCM.01274-10

Martin, D. P., Lemey, P., and Posada, D. (2011). Analysing recombination in nucleotide sequences. Mol. Ecol. Resour. 11, 943–955. doi:10.1111/j.1755-0998.2011.03026.x

Martin, D. P., Murrell, B., Golden, M., Khoosal, A., and Muhire, B. (2015). RDP4: Detection and analysis of recombination patterns in virus genomes. Virus Evol. 1:vev003. doi:10.1093/ve/vev003

Mawassi, M., Dror, O., Bar-Joseph, M., Piasezky, A., Sjölund, M. J., Levitzky, N., et al. (2018). ’Candidatus Liberibacter solanacearum’ is Tightly Associated with Carrot Yellows Symptoms in Israel and Transmitted by the Prevalent Psyllid Vector Bactericera trigonica. Phytopathology, “in press”. doi:10.1094/PHYTO-10-17-0348-R

Monger, W. A., and Jeffries, C. J. (2018). A survey of ‘*Candidatus* Liberibacter solanacearum’ in historical seed from collections of carrot and related Apiaceae species. Eur. J. Plant Pathol. 150, 803–815. doi:10.1007/s10658-017-1322-6

Munyaneza, J. E. (2015). Zebra chip disease, *Candidatus* liberibacter, and potato psyllid: A global threat to the potato industry. Am. J. Potato Res. 92, 230–235. doi:10.1007/s12230-015-9448-6

Munyaneza, J. E., Crosslin, J. M., and Upton, J. E. (2007). Association of *Bactericera cockerelli* (Homoptera: Psyllidae) with “zebra chip”, a new potato disease in southwestern United States and Mexico. J. Econ. Entomol. 100, 656–663. doi:10.1093/jee/100.3.656

Munyaneza, J. E., Fisher, T. W., Sengoda, V. G., Garczynski, S. F., Nissinen, A., and Lemmetty, A. (2010a). First report of ‘*Candidatus* Liberibacter solanacearum’ associated with psyllid-affected carrots in Europe. Plant Dis. 94:639. doi:10.1094/PDIS-94-5-0639A

Munyaneza, J. E., Fisher, T. W., Sengoda, V. G., Garczynski, S. F., Nissinen, A., and Lemmetty, A. (2010b). Association of “*Candidatus* Liberibacter solanacearum” with the psyllid *Trioza apicalis* (Hemiptera: Triozidae) in Europe. J. Econ. Entomol. 103, 1060–1070. doi:10.1603/EC10027

Munyaneza, J. E., Sengoda, V. G., Stegmark, R., Arvidsson, A. K., Anderbrant, O., Yuvaraj, J. K., et al. (2012a). First report of “*Candidatus* Liberibacter solanacearum” associated with psyllid-affected carrots in Sweden. Plant Dis. 96:453. doi:10.1094/PDIS-10-11-0871

Munyaneza, J. E., Sengoda, V. G., Sundheim, L., and Meadow, R. (2012b). First report of “*Candidatus* Liberibacter solanacearum” associated with psyllid-affected carrots in Norway. Plant Dis. 96:454. doi:10.1094/PDIS-10-11-0870

Munyaneza, J. E., Swisher, K. D., Hommes, M, Willhauck, A, Buck, H., and Meadow, R. (2015). First Report of ‘*Candidatus* Liberibacter solanacearum’ Associated With Psyllid-Infested Carrots in Germany. Plant Dis. 99:1269. doi:10.1094/PDIS-02-15-0206-PDN

Munyaneza, J. E., Mustafa, T., Fisher, T. W., Sengoda, V. G., and Horton, D. R. (2016) Assessing the likelihood of transmission of Candidatus Liberibacter solanacearum to carrot by potato psyllid, Bactericera cockerelli (Hemiptera: Triozidae). PLoS ONE 11:e0161016. doi:10.1371/journal.pone.0161016

Murray, M. G., and Thompson, W. F. (1980). Rapid isolation of high molecular weight plant DNA. Nucl. Acids Res. 8, 4321–4326. doi:10.1093/nar/8.19.4321

Nei, M., and Gojobori, T. (1986). Simple methods for estimating the numbers of synonymous and nonsynonymous nucleotide substitutions. Mol. Biol. Evol. 3, 418–426.

Nei, M. (1987). Molecular Evolutionary Genetics. New York, NY: Columbia University Press.

Nelson, W. R., Fisher, T. W., and Munyaneza, J. E. (2011). Haplotypes of “*Candidatus* Liberibacter solanacearum” suggest long-standing separation. Eur. J. Plant Pathol. 130, 5–12. doi:10.1007/s10658-010-9737-3

Nelson, W. R., Sengoda, V. G., Crosslin, J. M., Alfaro-Fernández, A. O., Font, M. I., and Munyaneza, J. E. (2013). A new haplotype of “*Candidatus* Liberibacter solanacearum” in the Mediterranean region. Eur. J. Plant Pathol. 135, 633–639. doi:10.1007/s10658-012-0121-3

Nissinen, A. I., Haapalainen, M., Jauhiainen, L., Lindman, M., and Pirhonen, M. (2014). Different symptoms in carrots caused by male and female carrot psyllid feeding and infection by ‘*Candidatus* Liberibacter solanacearum’. Plant Pathol. 63, 812–820. doi:10.1111/ppa.12144

Puttamuk, T., Zhou, L. J., Thaveechai, N., Zhang, S. A., Armstrong, C. M., and Duan, Y. P. (2014). Genetic diversity of *Candidatus* Liberibacter asiaticus based on two hypervariable effector genes in Thailand. PLoS ONE 9:e112968. doi:10.1371/journal.pone.0112968

Sarkar, S. F., and Guttman, D. S. (2004). Evolution of the core genome of *Pseudomonas syringae*, a highly clonal, endemic plant pathogen. Appl. Environ. Microbiol. 78, 1999–2012. doi: 0.1128/AEM.70.4.1999-2012.2004

Secor, G. A., Rivera-Varas, V., Abad, J. A., Lee, I. M., Clover, G. R. G., Liefting, L. W., et al. (2009). Association of ‘*Candidatus* Liberibacter solanacearum’ with zebra chip disease of potato established by graft and psyllid transmission, electron microscopy, and PCR. Plant Dis. 93, 574–583. doi:10.1094/PDIS-93-6-0574

Soliman, T., Mourits, M. C. M., Oude Lansink, A. G. J. M., and van der Werf, W. (2013). Economic justification for quarantine status – the case study of *Candidatus* Liberibacter Solanacearum in the European Union. Plant Pathol. 62, 1106–1113. doi:10.1111/ppa.12026

Swisher Grimm, K.D., Garczynski, S.F. (2018). Identification of a new haplotype of ‘*Candidatus* Liberibacter solanacearum’ In Solanum tuberosum. Plant Dis. 10.1094/pdis-06-18-0937-RE

Tahzima, R., Maes, M., Achbani, E. H., Swisher, K. D., Munyaneza, J. E., and De Jonghe, K. (2014). First report of ‘*Candidatus* Liberibacter solanacearum’ on carrot in Africa. Plant Dis. 98:1426. doi:10.1094/PDIS-05-14-0509-PDN

Tahzima, R., Massart, S., Achbani, E. H., Munyaneza, J. E., and Ouvrard, D. (2017). First report of ‘*Candidatus* Liberibacter solanacearum’ associated with the psyllid *Bactericera trigonica* Hodkinson on carrots in northern Africa. Plant Dis. 101:242. doi:10.1094/PDIS-07-16-0964-PDN

Tajima, F. (1989). Statistical method for testing the neutral mutation hypothesis by DNA polymorphism. Genetics 123, 585–595.

Tancos, M. A., Lange, H. W., and Smart, C. D. (2015). Characterizing the genetic diversity of the Clavibacter michiganensis subsp. michiganensis population in New York. Phytopathology 105, 169–179. doi:10.1094/PHYTO-06-14-0178-R

Teresani, G. R., Bertolini, E., Alfaro-Fernández, A., Martínez, C., Tanaka, F. A. O., Kitajima, E. W., et al. (2014). Association of ‘*Candidatus* Liberibacter solanacearum’ with a vegetative disorder of celery in Spain and development of real-time PCR method for its detection. Phytopathology 104, 804–811. doi:10.1094/PHYTO-07-13-0182-R

Thompson, S. M., Johnson, C. P., Lu, A. Y., Frampton, R. A., Sullivan, K. L., Fiers, M. W. E. J., et al. (2015). Genomes of ‘*Candidatus* Liberibacter solanacearum’ haplotype A from New Zealand and the United States suggest significant genome plasticity in the species. Phytopathology 105, 863–871. doi:10.1094/PHYTO-12-14-0363-FI

Timilsina, S., Jibrin, M. O., Potnis, N., Minsavage, G. V., Kebede, M., Schwartz, A., et al. (2015). Multilocus sequence analysis of *xanthomonads* causing bacterial spot of tomato and pepper plants reveals strains generated by recombination among species and recent global spread of *Xanthomonas gardneri*. Appl. Environ. Microbiol. 81, 1520–1529. doi:10.1128/AEM.03000-14

Trantas, E. A., Sarris, P. F., Mpalantinaki, E. E., Pentari, M. G., Ververidis, F. N., and Goumas, D. E. (2013). A new genomovar of *Pseudomonas cichorii*, a causal agent of tomato pith necrosis. Eur. J. Plant Pathol.137, 477–493. doi:10.1007/s10658-013-0258-8

Wang, J., Haapalainen, M., Schott, T., Thompson, S. M., Smith, G. R., Nissinen, A. I., et al. (2017). Genomic sequence of ‘*Candidatus* Liberibacter solanacearum’ haplotype C and its comparison with haplotype A and B genomes. PLoS ONE 12:e0171531. doi:10.1371/journal.pone.0171531

Watterson, G. A. (1975). On the number of segregating sites in genetical models without recombination. Theor. Popul. Biol. 7, 256–276. doi:10.1016/0040-5809(75)90020-9

Wen, A., Mallik, I., Alvarado, V. Y., Pasche, J. S., Wang, X., Li, W., et al. (2009). Detection, distribution, and genetic variability of ‘*Candidatus* Liberibacter’ species associated with the zebra complex disease of potato in the North America. Plant Dis. 93, 1102–1115. doi:10.1094/PDIS-93-11-1102

Wu, F., Deng, X., Liang, G., Wallis, C., Trumble, J. T., Prager, S., et al. (2015). De Novo genome sequence of “Candidatus Liberibacter solanacearum” from a single potato psyllid in California. Genome Announc. 3:e01500–15. doi:10.1128/genomeA.01500-15

Young, J. M., Park, D. C., Shearman, H. M., and Fargier, E. (2008). A multilocus sequence analysis of the genus *Xanthomonas*. Syst. Appl. Microbiol. 31, 366–377. doi:10.1016/j.syapm.2008.06.004

Zheng, Z., Bao, M., Wu, F., Chen, J., and Deng, X. (2016). Predominance of single prophage carrying a CRISPR/*cas* system in *“Candidatus* Liberibacter asiaticus*”* strains in southern China. PLoS ONE 11:e0146422. doi:10.1371/journal.pone.0146422

Zheng, Z., Bao, M., Wu, F., Van Horn, C., Chen, J., and Deng, X. (2018). A Type 3 Prophage of ‘*Candidatus* Liberibacter asiaticus’ Carrying a Restriction-Modification System. Phytopathology, 100, 454–461. doi:10.1094/PHYTO-08-17-0282-R

Zhou, L., Powell, C., Li, W., and Duan, Y. (2013). Prophage-mediated dynamics of ‘*Candidatus* Liberibacter asiaticus’ populations, the destructive bacterial pathogens in citrus huanglongbing. PLoS ONE 8:e82248. doi:10.1371/journal.pone.0082248

