## Supplementary material for "New Insights into the Genetic Diversity of the Bacterial Plant Pathogen ‘*Candidatus* Liberibacter solanacearum’ as Revealed by a New Multilocus Sequence Analysis Scheme"

**Abstract**

'*Candidatus* Liberibacter solanacearum' (Lso) has emerged as a serious threat on solanaceous and apiaceous crops worldwide. Five Lso haplotypes (LsoA, LsoB, LsoC, LsoD and LsoE) have been identified so far. To decipher genetic relationships between Lso strains, a MLSA study of seven housekeeping genes (*acnA*, *atpD*, *ftsZ*, *glnA*, *glyA*, *gnd* and *groEL*) was performed on a representative bacterial collection of 49 Lso strains. In all, 5415 bp spanning the seven loci were obtained from each of the 49 strains of our bacterial collection. Analysis of sequence data was consistent with a clonal population structure with no evidence of recombination. Phylogenies reconstructed from individual genes, and with concatenated data, were globally congruent with each other. In addition to the five highly supported and distinct genetic clusters, which correspond to the five established haplotypes, our phylogenetic data revealed the presence of a sixth haplotype, designated 'LsoG'. This new haplotype is currently represented by two strains from France which had distinct sequences in four out of the seven tested housekeeping genes. Altogether, the data presented here provide new information regarding the genetic structure of Lso and the evolutionary history of the haplotypes defined within this bacterial species.

**Running title:** '*Candidatus* Liberibacter solanacearum' genetic diversity

**Keywords:** '*Candidatus* Liberibacter solanacearum', genetic diversity, haplotype, MLSA, apiaceous and solanaceous crop

### Supplementary material

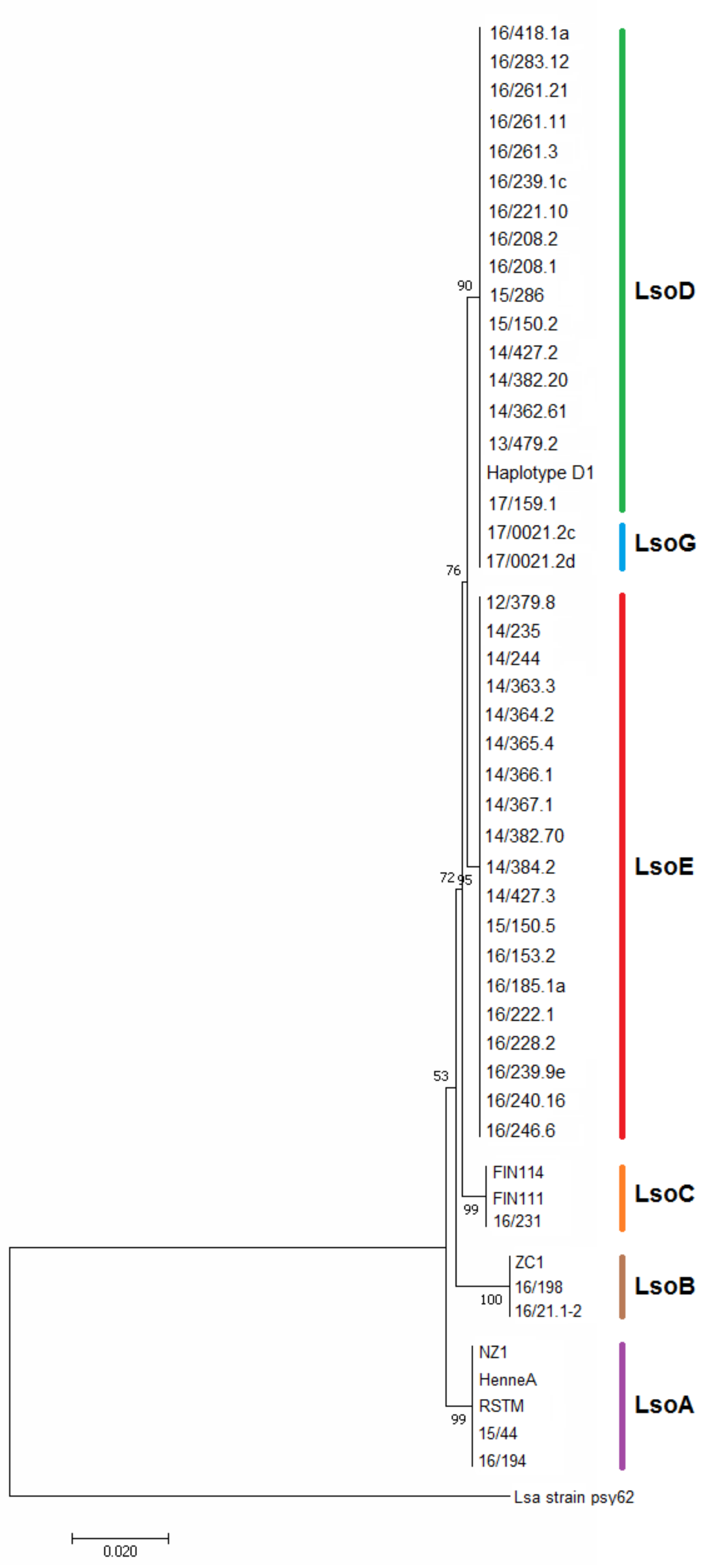


**Figure S1.** Phylogenetic tree based on the *acnA* gene among the 49 Lso strains. The tree was constructed using the maximum likelihood method. Boostrap values over 50 (1,000 replicates) were shown at each node. The scale bar indicates the number of nucleotide substitutions per site. The tree is rooted with the '*Ca.* L. asiaticus' strain psy62.

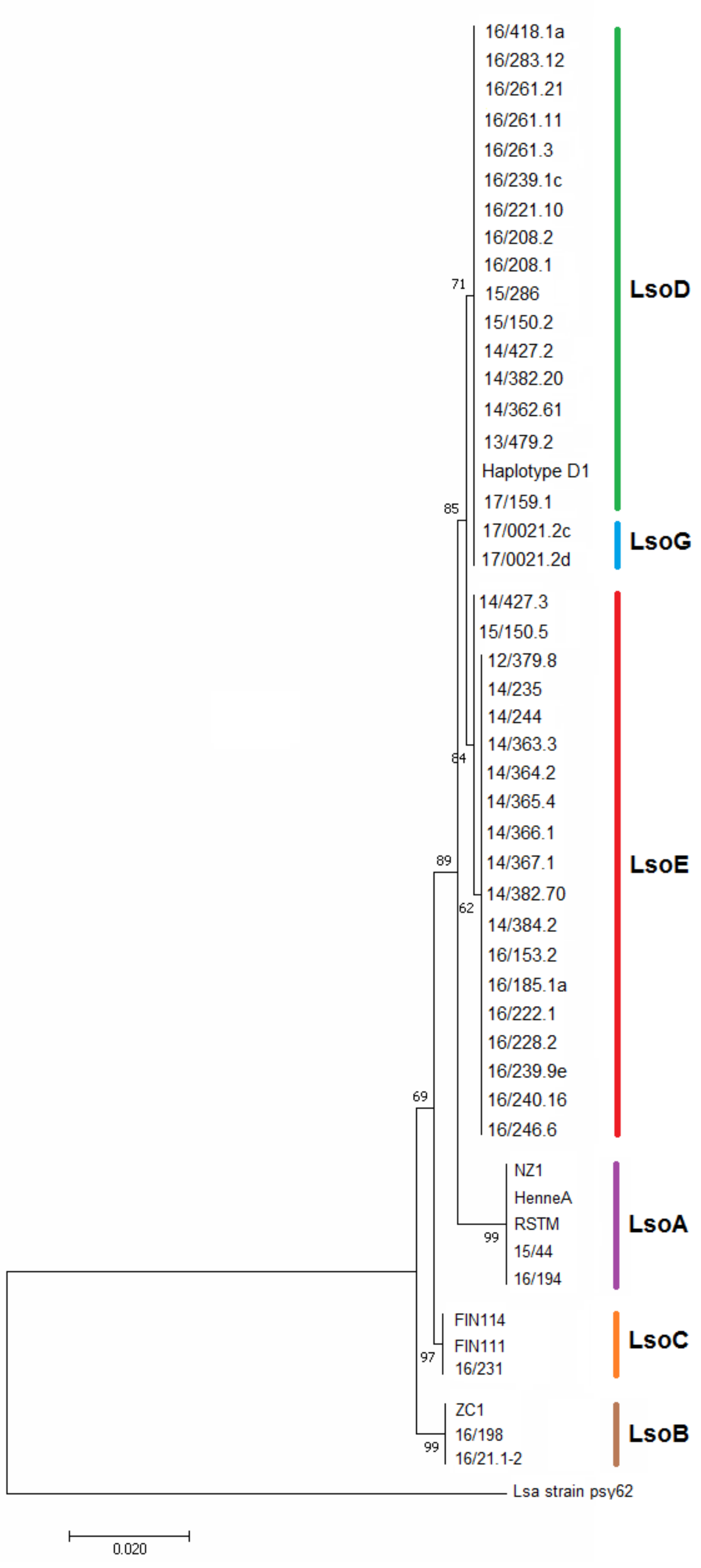


**FIGURE S2.** Phylogenetic tree based on the *atpD* gene among the 49 Lso strains. The tree was constructed using the maximum likelihood method. Bootstrap values over 50 (1,000 replicates) were shown at each node. the scale bar indicates the number of nucleotide substitutions per site. The tree is rooted with the '*Ca*. L. asiaticus strain psy62

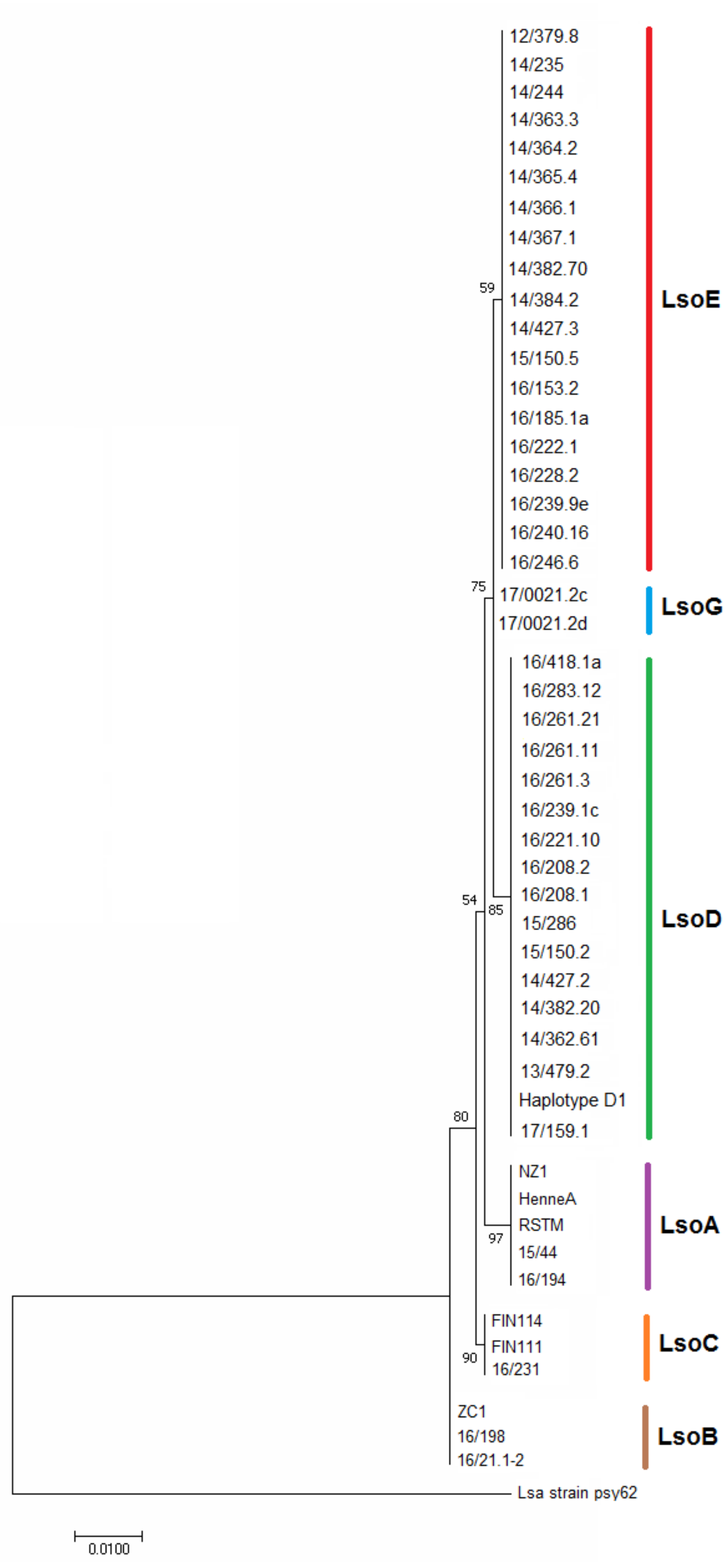


**FIGURE S3.** Phylogenetic tree based on the *ftsZ* gene among the 49 Lso strains. The tree was constructed using the maximum likelihood method. Boostrap values over 50 (1,000 replicates) were shown at each node. The scale bar indicates the number of nucleotides substitutions per site. The tree is rooted with the '*Ca.* L. asiaticus' strain psy62.

39

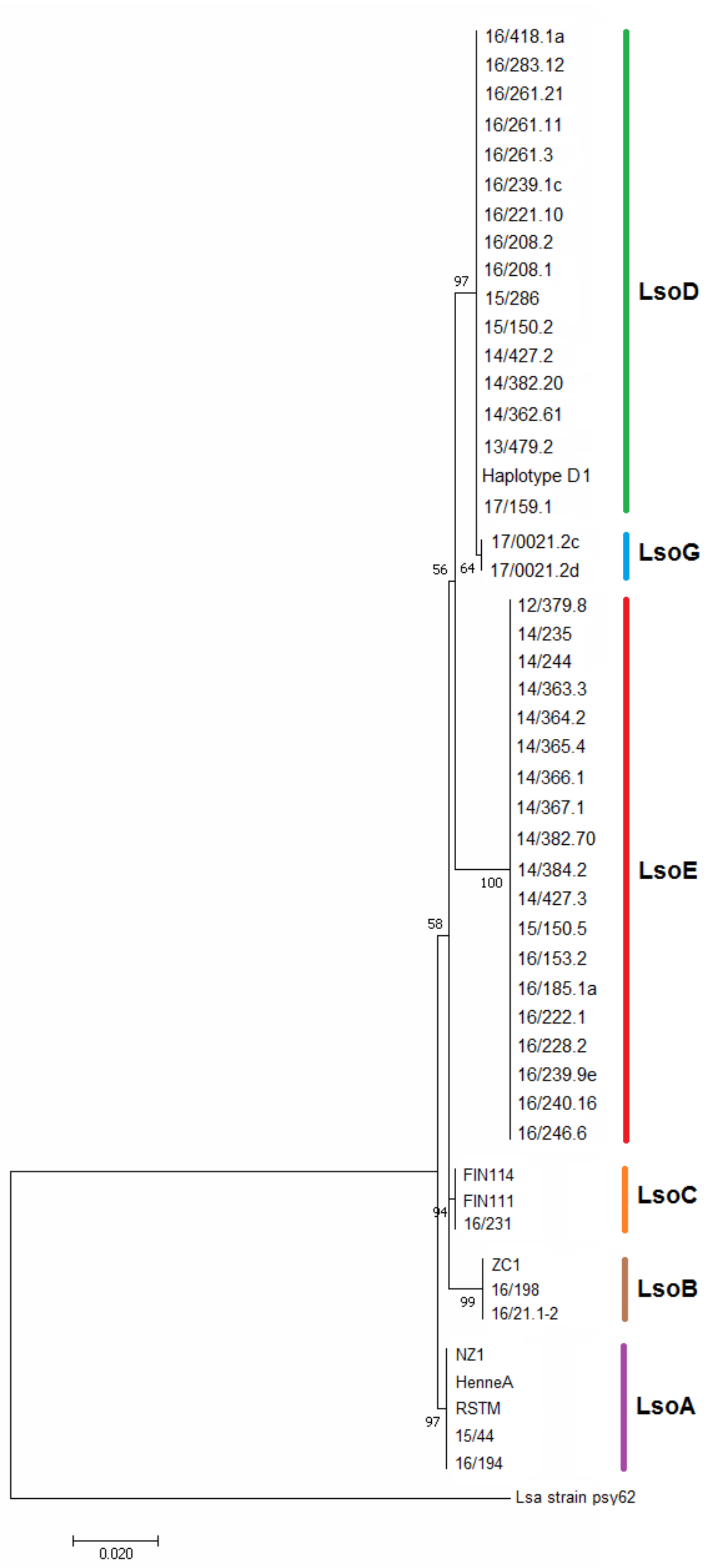


**FIGURE S4.** Phylogenetic tree based on the *glnA* gene among the 49 Lso strains. The tree was constructed using the maximum likelihood method. Boostrap values over 50 (1,000 replicates) were shown at each node. The scale bar indicates the number of nucleotide substitutions per site. The tree is rooted with the '*Ca.* L. asiaticus' strain psy62.

40

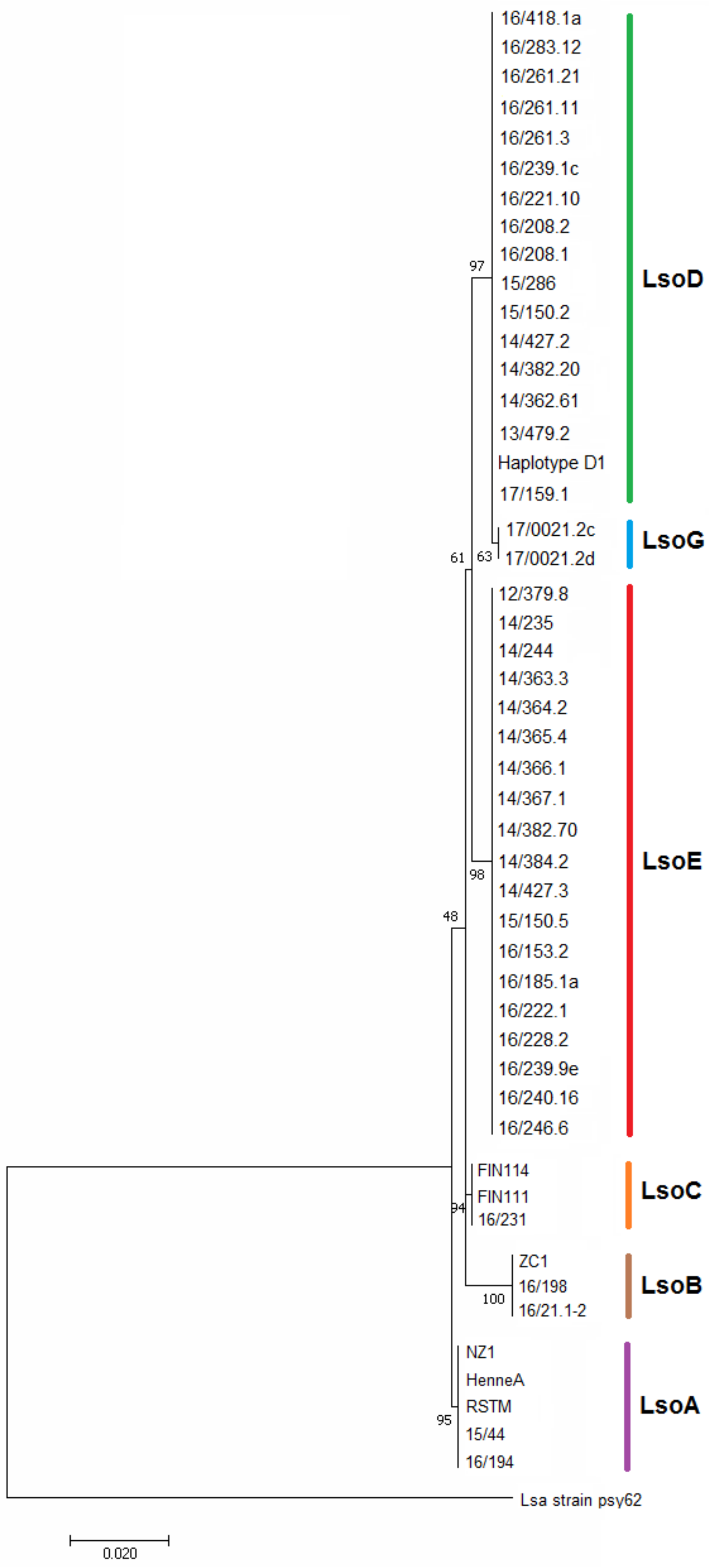


**FIGURE S5.** Phylogenetic tree based on the *glyA* gene among the 49 Lso strains. The tree was constructed using the maximum likelihood method. Boostrap values over 50 (1,000 replicates) were shown at each node. The scale bar indicates the number of nucleotide substitutions per site. The tree is rooted with the '*Ca.* L. asiaticus' strain psy62.

41

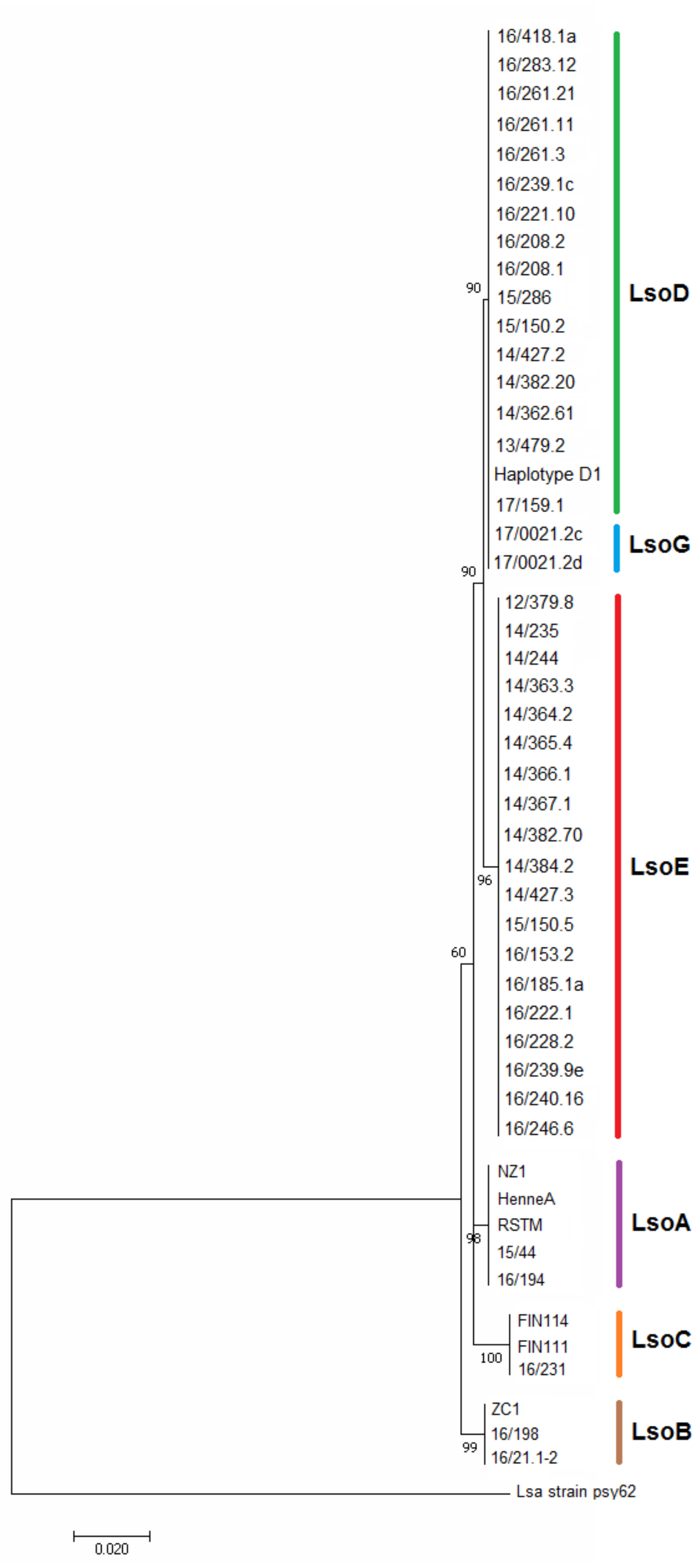


**FIGURE S6.** Phylogenetic tree based on the *gnd* gene among the 49 Lso strains. The tree was constructed using the maximum likelihood method. Boostrap values over 50 (1,000 replicates) were shown at each node. The scale bar indicates the number of nucleotide substitutions per site. the tree is rooted with the '*Ca*. L. asiaticus' strain psy62.

42

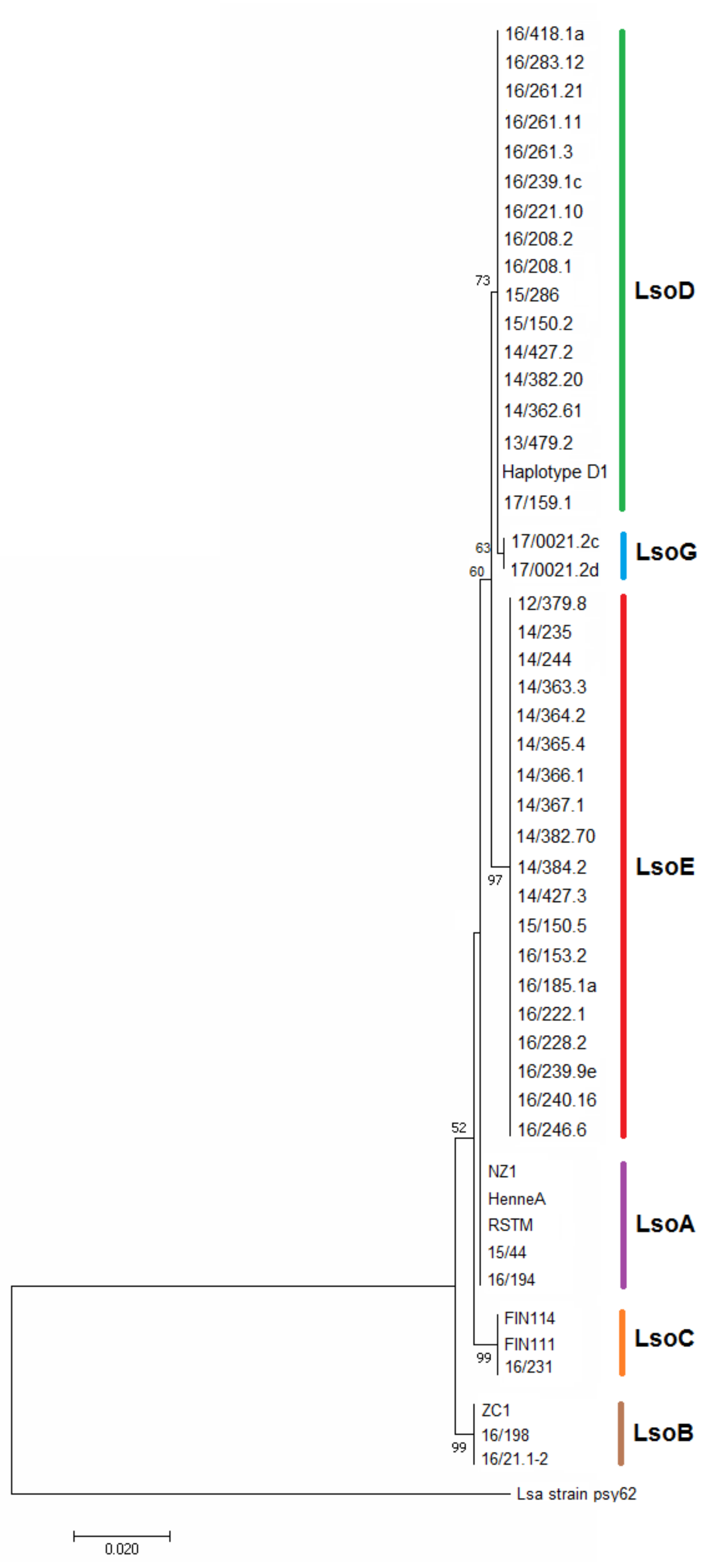


**FIGURE S7.** Phylogenetic tree based on the *groEL* gene among the 49 Lso strains. The tree was constructed using the maximum likelihood method. Boostrap values over 50 (1,000 replicates) were shown at each node. The scale bar indicates the number of nucleotide subsititutions per site. The tree is rooted with the '*Ca.* L. asiaticus' strain psy62.

### Tables

**Table 1.** List of Lso strains used in this study

| Sample ID | Source of isolation | Geographic origin | Year of isolation | Lso haplotype[1] |
|---|---|---|---|---|
| NZ1 | *Bactericera cockerelli* | New Zealand | 2010 | |
| RSTM | *Bactericera cockerelli* | United States | 2015 | |
| 16/19.4 | *Bactericera cockerelli* | United States | NA | A |
| HenneA | *Bactericera cockerelli* | USA | 2012 | |
| 15/44 | *Solanum lycopersicum* | New Zealand | 2010 | |
| 16/19.8 | *Bactericera cockerelli* | United States | NA | |
| ZC1 | *Bactericera cockerelli* | United States | NA | B |
| 16/21.1-2 | *Solanum tuberosum* | United States | NA | |
| FIN111 | *Trioza apicalis* | Finland | 2012 | |
| FIN114 | *Trioza apicalis* | Finland | 2012 | C |
| 16/231 | *Trioza apicalis* | Finland | NA | |
| 16/239-1c | *Bactericera trigonica* | Eure-et-Loir, France | 2016 | |
| Haplotype D1 | *Bactericera trigonica* | Israel | 2017 | |
| 15/150.2 | *Daucus carota* | Canary Islands, Spain | NA | |
| 13/479.2 | *Daucus carota* | Dordogne, France | 2013 | |
| 16/418.1a | *Daucus carota* | Dordogne, France | 2016 | |
| 16/221-10 | *Daucus carota* | Eure-et-Loir, France | 2016 | |
| 14/427.2 | *Daucus carota* | Gard, France | 2014 | |
| 16/208-2 | *Daucus carota* | Gers, France | 2016 | |
| 16/261-21 | *Daucus carota* | Haute-garonne, France | 2016 | D |
| 16/283-12 | *Daucus carota* | Loiret, France | 2016 | |
| 16/208-1 | *Daucus carota* | Lot-et-Garonne, France | 2016 | |
| 14/362.61 | *Daucus carota* | Maine et Loire, France | 2014 | |
| 14/382.20 | *Daucus carota* | Maine et Loire, France | 2014 | |
| 17/159.1 | *Daucus carota* | Morocco | 2013 | |
| 16/261-3 | *Daucus carota* | Tarn, France | 2016 | |
| 16/261-11 | *Daucus carota* | Tarn, France | 2016 | |
| 15/286 | *Petroselinum crispum* | Aude, France | 2015 | |
| 14/367.1 | *Anthriscus cerefolium* | Maine et Loire, France | 2014 | |
| 14/363.3 | *Apium graveolens* | Maine et Loire, France | 2014 | |
| 16/239-9e | *Bactericera trigonica* | Loiret, France | 2016 | |
| 16/153.2 | *Bactericera trigonica* | Segovia, Spain | 2014 | |
| 15/150.5 | *Daucus carota* | Canary Islands, Spain | NA | |
| 16/240-16 | *Daucus carota* | Cher, France | 2016 | |
| 14/244 | *Daucus carota* | Eure-et-Loir, France | 2014 | |
| 14/427.3 | *Daucus carota* | Gard, France | 2014 | |
| 14/235 | *Daucus carota* | Loiret, France | 2014 | |
| 12/379.8 | *Daucus carota* | Loir-et-Cher, France | 2012 | E |
| 16/185-1a | *Daucus carota* | Loir-et-Cher, France | 2016 | |
| 14/382.70 | *Daucus carota* | Maine et Loire, France | 2014 | |
| 16/246-6 | *Daucus carota* | Maine et Loire, France | 2016 | |
| 16/228-2 | *Daucus carota* | Vienne, France | 2016 | |
| 14/366.1 | *Foeniculum vulgare* | Maine et Loire, France | 2014 | |
| 14/365.4 | *Pastinaca sativa* | Maine et Loire, France | 2014 | |
| 16/222-1 | *Petroselinum crispum* | Eure-et-Loir, France | 2016 | |
| 14/364.2 | *Petroselinum crispum* | Maine et Loire, France | 2014 | |
| 14/384.2 | *Petroselinum crispum* | Maine et Loire, France | 2014 | |
| 17/0021-2c | *Daucus carota* | Marne, France | 2017 | G |
| 17/0021-2d | *Daucus carota* | Marne, France | 2017 | |

[1]: Lso haplotype was determined during previous study (Hajri et al., 2017) using Nelson et al. (2011) or during this study by building the phylogenetic trees including strains LsoA (strains NZ1, HenneA and RSTM), LsoB (strain ZC1), LsoC (strains FIN111 and FIN114) and LsoD (strain haplotype D1); NA, not available

**Table 2.** Primers of housekeeping genes used for amplification and sequencing

| Locus | Primer name | Sequence (5'-3') | Size of amplicon (bp) |
|---|---|---|---|
| *acnA* | acnA-F | AATCTTGTTGGTTTTGGATGTACAACGTG | 949 |
| | acnA-R | CTATCAAATTGGAGCGATGTATACGCT | |
| *atpD* | atpD-F | GCGGGAGTTGGGAAAACAGTATTAATCAT | 811 |
| | atpD-R | ATTCAGCAACATGAAACGGCTGTGACAT | |
| *glnA* | glnA-F | ATGGTTGATGACGCTACCTCTATCATC | 905 |
| | glnA-R | AGGGCTTTCGCATGTTTGATAATACCTC | |
| *glyA* | glyA-F | CGTTCAACCTCATTCTGGATCTCAGATGA | 843 |
| | glyA-R | AATCCTCTCGTTGTACCAGATGGCGTA | |
| *gnd* | gnd-F | CTAATGATGATAACAGACGGGAATCC | 889 |
| | gnd-R | GTATTATACATCCTGCACGCCAGA | |
| *groEL* | groEL-F | CAATCTCGCTGTTCAAGAAGTTGTAGA | 927 |
| | groEL-R | TTAACAGATAATGCTTGAGCAGCTCG | |
| *ftsZ* | ftsZ-F | ATGGTGGAAAAACACTCTAATGTGGATAT | 915 |
| | ftsZ-R | AGCCTCATCAAATGTAGCACCAAGAAT | |

**Table 3.** Summary statistics for the seven housekeeping genes and concatenated sequences used in this study

| Locus | sites[a] | GC% | S[b] | $\theta\pi$[c] | $\theta w$[d] | Tajima's D[e] | d*N*/d*S*[f] |
|---|---|---|---|---|---|---|---|
| *acnA* | 816 | 37.2 | 25 | 0.00640 | 0.00687 | -0.22568 | 0.128 |
| *atpD* | 759 | 40.7 | 20 | 0.00578 | 0.00591 | -0.06851 | 0.030 |
| *ftsZ* | 774 | 43.2 | 12 | 0.00390 | 0.00348 | 0.35732 | 0.021 |
| *glnA* | 783 | 35.5 | 27 | 0.01123 | 0.00773 | 1.49289 | 0.311 |
| *glyA* | 747 | 41.8 | 19 | 0.00648 | 0.00570 | 0.43441 | 0.199 |
| *gnd* | 744 | 33.9 | 22 | 0.00642 | 0.00693 | -0.24289 | 0.760 |
| *groEL* | 792 | 39.9 | 18 | 0.00521 | 0.00510 | 0.06685 | 0.000 |
| Concat[g] | 5415 | 38.9 | 143 | 0.00649 | 0.00597 | 0.31854 | 0.161 |

[a] number of analyzed sites
[b] number of polymorphic sites
[c] nucleotide diversity (Nei, 1987)
[d] nucleotide diversity with Watterson's estimator (Watterson, 1975)
[e] Neutrality test of Tajima (1989); all values are not significant ($P > 0.10$)
[f] ratio of non-synonymous to synonymous substitutions
[g] data set of concatenated sequences of the seven loci

**Table 4.** New SNP features of the LsoG haplotype

| | Haplotypes | | | | | |
|---|---|---|---|---|---|---|
| **Gene region** | **A** | **B** | **C** | **D** | **E** | **G** |
| *glnA* (778) | G | G | G | G | G | A |
| *glyA* (393) | C | C | C | C | C | G |
| *groEL* (726) | G | G | G | G | G | T |
| *ftsZ* (708) | C | C | C | T | C | C |
| *ftsZ* (717) | G | G | G | A | G | G |

Nucleotide numbers are indicated relative to the first nucleotide of the aligned sequence of each gene
